# MetA is a ‘thermal fuse’ that arrests growth and protects *Escherichia coli* at elevated temperatures

**DOI:** 10.1101/2021.06.14.448417

**Authors:** Severin Schink, Zara Gough, Elena Biselli, Mariel Garcia Huiman, Yu-Fang Chang, Markus Basan, Ulrich Gerland

**Author notes:** Contributed equally (Severin Schink, Zara Gough, Elena Biselli).

## Abstract

Adaptive stress resistance in microbes is mostly attributed to the expression of stress response genes, such as heat shock proteins, which prevent deterioration of cellular material. Here, we report a novel response of *E. coli* to heat stress: induction of a growth-arrested state, caused by degradation of an enzyme in the methionine biosynthesis pathway (MetA). While MetA degradation is detrimental for proliferation, we show that the resulting growth arrest has a direct benefit for survival at high temperatures; it protects cells when temperatures rise beyond 50°C, increasing the survival chances by over 1000-fold. Using a combination of experiments and mathematical modelling, we show that degradation of MetA leads to the coexistence of growing and non-growing cells, allowing microbes to bet-hedge between continued growth if conditions remain bearable and survival if conditions worsen. We test our model experimentally and verify quantitatively how protein expression, degradation rates and environmental stresses affect the partitioning between growing and non-growing cells. Because growth arrest can be abolished with simple mutations, such as point mutations of MetA and knock-outs of proteases, we interpret the breakdown of methionine synthesis as a system that has evolved to disintegrate at high temperature and shut off growth, analogous to ‘thermal fuses’ used in engineering to shut off electricity when the device could be damaged by overheating.

## Introduction

Microbes in nature face a variety of environments, ranging from ideal growth conditions to hostile environments, where stressors threaten their survival. One such stressor is elevated temperature, which can arise quickly and unpredictably. When temperatures rise above the optimal range, microbes employ a well characterized heat shock response ^1–3^. Molecular chaperones protect proteins in the cytoplasm and periplasm by maintaining their native conformation ^4,1,2^ and proteases degrade misfolded and aggregated proteins ^5^. DNA repair is upregulated^6^ and toxic reactive oxygen species, which are at particularly high levels at elevated temperatures ^7^, are scavenged by catalases ^8^, superoxide dismutases ^9^, and small molecules like glutathione ^10,11^. The induction of these protective genes, collectively referred to as the heat shock response, permits cells to maintain their functionality and recover once conditions have improved.

However, some responses of microbes appear less logical. A long-standing puzzle is the active degradation of a key enzyme in the methionine biosynthesis pathway, homoserine O-succinyltransferase (MetA), when temperatures increase ^12–14^. This degradation causes methionine limitation, which slows the growth of cultures not supplemented with methionine ^12,15^ and decreases the maximum growth temperature in minimal medium from 47°C to 45°C ^16^. The degradation of MetA was found across bacterial species from Enterobacteriaceae, including *Escherichia coli, Salmonella typhimurium* and *Klebsiella aerogenes*^17^ to Alphaproteobacteria like *Agrobacterium fabrum* ^18^. Even in the gram-positive *Paenibacillus polymyxa*^19^ and *Corynebacterium glutamicum*^20^, in which MetA and MetXA, respectively, encode a homoserine o-acetyltransferase rather than a o-succinyltransferase, the temperature-dependent degradation is conserved. This conservation is conspicuous because MetA is only weakly conserved between these species (Fig. S1), but is the only biosynthetic enzyme which is degraded at high rates ^15^. Furthermore, MetA degradation is not inevitable, since several point-mutations were discovered that decrease degradation rates, alleviate growth defects and increase the maximum growth temperature ^16,21^. Ron and coworkers hypothesized that MetA could serve as a regulatory protein ^14,15,17^, given that it displays rapid ATP-dependent proteolysis in combination with activation of transcription ^13,14^, a hallmark of regulatory proteins that enable quick response times ^22^. However, it is unclear why stalling methionine biosynthesis should be actively regulated by bacteria, because there would need to be “some undisclosed selective benefit”, as Ron and Davies put it ^12^. To date, the only described benefit is an increase in antibiotic persistence at elevated temperatures, discovered by Mordukhova and Pan ^23^, but there is no apparent relation between antibiotic stress and heat stress.

Here, we identify the missing selective benefit of MetA degradation. Furthermore, we confirm the hypothesis that MetA dual-serves as a biosynthetic protein and a temperature-sensitive regulatory protein, and clarify the mechanism whereby MetA controls growth at elevated temperatures. Using *Escherichia coli* as a model system, we show that MetA plays the central role in a new type of regulated growth arrest that protects from cell death when temperature is further increased. The protective effect of growth arrest against heat shocks appears to be a more general physiological feature of microbes, as we can reproduce the phenomenon in *Saccharomyces cerevisiae*. Hence, heat-induced growth arrest appears to implement a ‘heat persistence’ strategy that is conceptually similar to antibiotic persistence ^24^. While the regulatory mechanism controlling antibiotic persistence remains elusive, we find that heat persistence is caused by a bistability in growth that is modulated by MetA degradation.

## RESULTS

### Elevated temperatures induce growth arrest in E. coli

To probe how heat affects the physiology of bacteria, we exposed cultures of *Escherichia coli* K-12 grown in minimal medium at elevated temperatures to a period of carbon starvation. During this period, the absence of *de novo* synthesis reveals the deteriorative effects of temperature on the cell. After the starvation period, we reinoculated the culture in fresh medium with a carbon substrate and observed the resulting recovery of the stressed culture. The temperature was kept constant throughout the entire experiment, i.e. initial growth, starvation, and recovery. We observed that the regrowth during the recovery phase displays lag times that depend on temperature (Fig. 1A). At 37°C, cultures recovered quickly, while at 45°C, the maximum growth temperature, the culture required more than 10 h to recover. Such long recovery times at high temperatures are puzzling, because the majority of bacteria were still viable at the time of reinoculation, as assayed by plating on LB agar at 37°C (Fig. S2), implying that the long lag times are not due to cell death, but instead caused by viable bacteria not growing.

**Figure 1.**
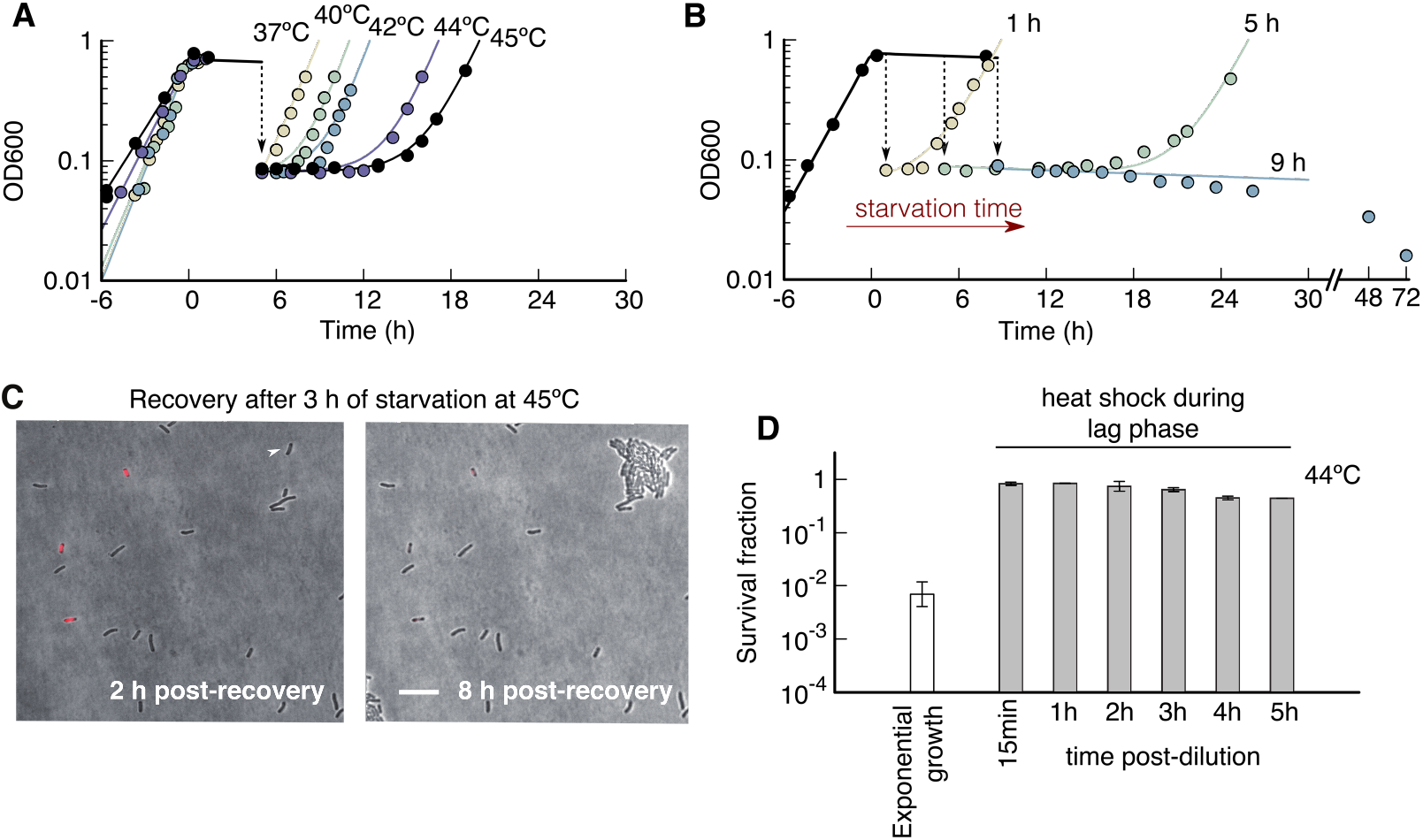
Growth arrest at high temperatures protects Escherichia coli in heat shocks. ***(A)** Optical density OD*_600_ *of Escherichia coli K-12 grown until nutrient depletion, first starved for 5 hours, then diluted into fresh medium to allow recovery. Colors denote culture temperatures, kept constant throughout each experiment. Lag times of cultures increase with temperature, from ~23 min at 37*°C, *to 11.1 h at 45*°C. *In panels A-C, growth is fitted as exponential N*(*t*) = *N*(0)*e^μt^ and recovery as a combination of decay and growth N*(*t*) = *N*_1_*e^−γt^* + *N*_2_*e^μt^*, *with γ* = 0.1 *h*^−1^. *Lag times are extracted as detailed in Methods. **(B)** Lag times at 45°C increase with starvation duration. No recovery was recorded within 3 days for 9 h starvation (blue) or longer. **(C)** Snapshots of time-lapse movies during recovery at 45*°C *on glycerol minimal medium agar pads, see Methods. Fluorescence of propidium iodide (red) reports permeability. One growing cell (white arrow) grows to a microcolony after 8 h of incubation. Scale bar: 10 μm. (**D**) Killing by heat shock (52°C, 10min) of E. coli cultured at 44°C in exponential growth (white) or during lag phase after 3 h of starvation and dilution (grey). E. coli growth arrested during lag phase are completely protected from heat killing at 52°C*.

The lag time also increases with the prior starvation duration (Fig. 1B), suggesting that cells gradually lose the ability to grow. For 1 hour of starvation, cultures recovered within 2 hours, but if starvation took longer than 9 h at 45°C, cultures did not recover within the 3 days of the experiment.

There are two qualitatively different scenarios for how long lag times can arise. One scenario is a homogeneous population of cells that all require a similarly long time to recover. The other scenario has two subpopulations, a large non-growing population and a small growing population. In the second case, the lag time is given by the time it takes for the small, growing population to dominate the cell count. To distinguish between these scenarios, we assayed the recovery phase at the single cell level: we extracted samples of a starved culture at 45°C and deposited them on pre-warmed agar pads supplemented with carbon substrate to allow recovery. We stained the bacteria with propidium iodide (red) for membrane permeability to detect cell death, and monitored growth by time-lapse microscopy (see Methods). Figure 1C shows two snapshots from a time-lapse, 2 h and 8 h after nutrient addition: The culture, previously starved for 3 h, shows three distinct populations. 5% of the population stains positive for cell permeability and are likely dead. A second, small subpopulation, 2%, recovered after about 2 hours, and after 8 h has formed micro-colonies, which are visible in Figure 1C (white arrow). The third and major subpopulation, 93%, stained negative for cell permeability, but did not grow within the 12 h of this time lapse. This data shows that recovery is highly heterogeneous; few cells recover early and dominate the recovery.

In an experimental scenario similar to ours, Trip & Youk recently observed that *Saccharomyces cerevisiae* arrest growth when cultures are diluted during exponential growth at elevated temperatures^11^. Indeed, this density-dependent growth behavior is shared by *E. coli*: we found that if cultures are diluted during exponential growth, they also enter growth arrest (Fig. S5A). This poses an interesting possibility that heat-induced growth arrest is a more general hallmark of microbes, and triggers the question: Is there a fitness benefit to heat-induced growth arrest ?

### Growth-arrest protects E. coli from heat shocks

Because growth arrest is unique to temperatures close to the growth limit (40-45°C for *E. coli*), but in a regime where conditions still allow growth, we wondered if growth arrest is an evolutionarily ‘chosen’ strategy, rather than an accident. For such a strategy to evolve, there would need to be a selective benefit associated with growth arrest, which outweighs the cost of reduced growth. To examine possible selective benefits, we exposed *E. coli* cultures to sudden heat shocks. We used our standard starvation protocol (Fig. 1A-C) to induce growth arrest, using a 5h period of starvation and subsequent reinoculation in fresh medium at 44°C. We took samples from the culture every hour after reinoculation, heat shocked them for 10 min at 52°C, and measured the resulting viability. Strikingly, growth-arrested cultures survived heat shocks almost unscathed, while in cultures heat shocked in exponential growth, only about 0.7% survived (Fig. 1D). The existence of this benefit shows that growth arrest could in principle be used by microorganisms as a part of the response to heat stress. To investigate this possibilty, we next sought to understand what causes *E. coli* to enter growth arrest.

### Breakdown of methionine synthesis causes growth arrest in E. coli

It is known that high temperature leads to denaturation of the proteome ^25,26^. One protein, homoserine O-succinyltransferase (MetA), which catalyzes the first commitment step of the methionine biosynthesis pathway, is particularly susceptible to heat stress ^12–14^. This enzyme aggregates and is degraded ^15^. To test if the long lag times are due to methionine limitation, we supplemented the medium with methionine and found a drastic reduction in lag time, from over 10 hours without methionine, down to less than 30 minutes with methionine (Fig. 2A). Similarly, growth arrest induced by dilution was also alleviated by the presence of methionine, decreasing the minimum initial population density from which cultures could regrow 100-fold (Fig S5). Investigating lag times on a single-cell level, we found that if the agar pad is supplemented with methionine, the majority of the cells recovered after starvation (63% compared to 2% w/o methionine), while the growth-arrested and non-permeable subpopulation virtually disappeared (5% compared to 93% w/o methionine), see Figure 2B. The absence of the growth-arrested subpopoulation (Fig. 2B) is consistent with the drastic reduction in lag times observed in bulk measurements (Fig. 2A).

**Figure 2.**
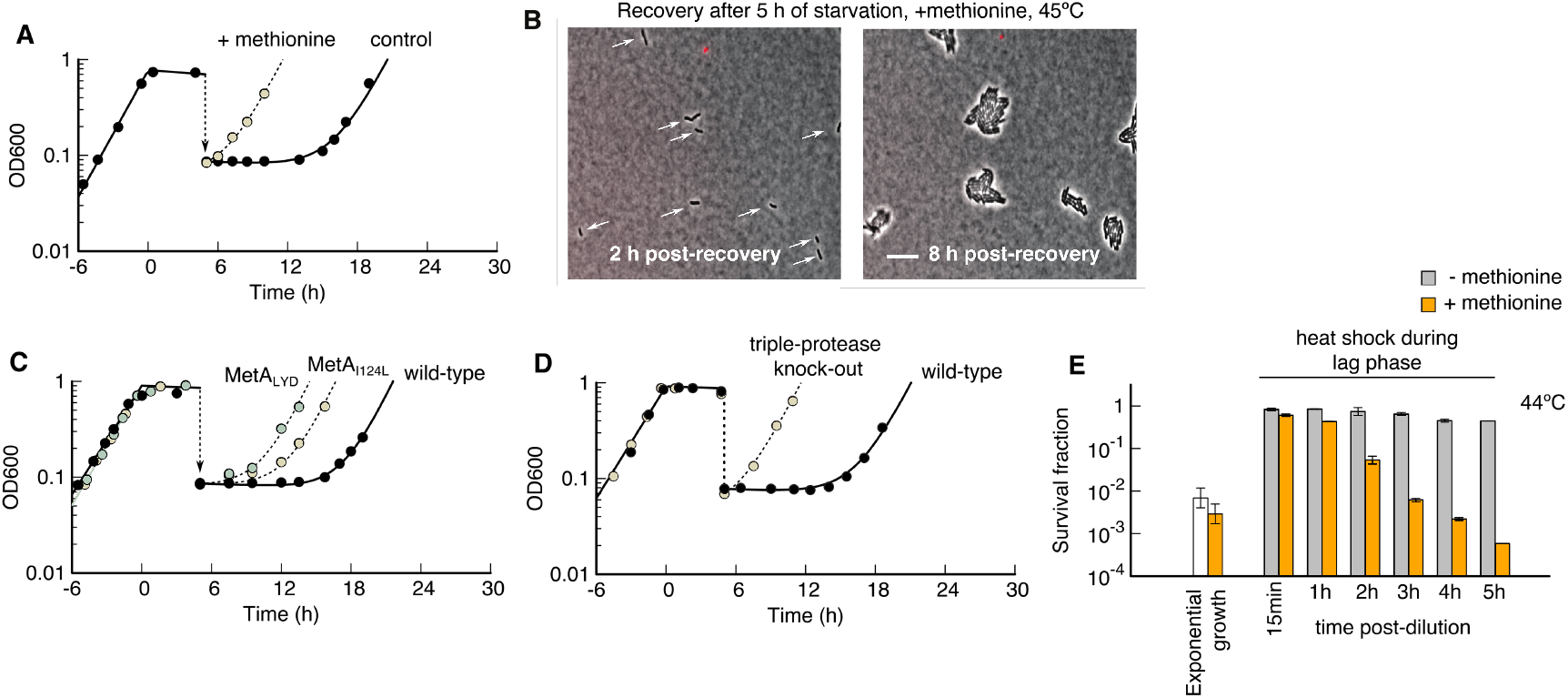
Growth arrested cells are methionine limited due to MetA degradation. ***(A)** Cultures supplemented with methionine recover quickly. Lag times (45°C, 5 h starvation) in cultures supplemented with methionine during the entire experiment are more than 90% shorter than without methionine* (*T_lag_* = (0.64 ± 0.02) *h compared to T_lag_* = (11.08 ± 0.4) *h, N = 2, p < 0.001). (**B**) Snapshots of a time-lapse of recovery on glycerol agar pads supplemented with methionine. Cells indicated with white arrows recover and form microcolonies after 8 hours. Scale bar: 10μm. (**C**) Comparison of wild-type MetA with two thermostable mutants. Lag times (45°C, 5 h starvation) decrease in mutants MetA I124Y (T_lag_* = (5.2 ± 1.0) *h, N = 3, p < 0.001) and LYD (T_lag_* = (3.7 ± 0.9) *h, N = 3, p < 0.001) compared to wild-type (T_lag_* = (11.1 ± 0.3) *h, N = 3). (**D**) Comparison of wild-type E. coli with a triple protease knock-out (Δlon, ΔhslV & ΔclpP). Absence of protease activity decreases lag time* (*T_lag_* = (1.1 ± 0.3) *h, N = 3, p = 0.0012) compared to wild-type (T_lag_* = (10.4 ± 1.6) *h, N = 3). (**E**) Heat shocks (10 min at 52°C) during exponential growth and during lag times at 44°C in cultures supplemented with and without methionine. Killing of the methionine-supplemented culture coincides with growth (at 2h: 2-fold increase of CFU/ml, at 5h: 20-fold increase of CFU/ml)*.

To test if methionine limitation is due to MetA degradation, and not a different enzyme in the methionine pathway, we tested MetA mutants with reduced degradation rates, either due to a single point mutation (I124L) or three point mutations (I124L-I229Y-N267D – abbr. LYD) identified by Mordukhova and coworkers ^21^. Indeed, these mutants, which have 2-3 fold decreased degradation rates ^21^, showed a significant decrease in lag times from about 11 h to about 5 h for the single point mutant (I124L) and 4 h for the triple-point mutant (LYD) (Fig. 2C).

When we hypothesize growth arrest to be a response to heat stress, an important question is whether MetA is degraded because it is broken, e.g. aggregated or otherwise damaged, or whether MetA is degraded despite being still functional. The latter option would speak for growth arrest being a ‘self-inflicted injury’, rather than an accident. To distinguish these two possibilities, we constructed a triple knock-out of proteases Δ*lon, ΔhslV* and Δ*clpP*, incapable of degrading cytoplasmic proteins, in particular MetA^14^. If MetA is broken, lag times should either remain unchanged or increase in this triple protease knock-out mutant. On the other hand, if functional MetA is actively degraded in wild-type, lag times will decrease in the mutant. In the experiment, we found a striking decrease of lag time from 10 h in wild-type down to 1 h in the triple protease knock-out (Fig. 2D), strongly suggesting that methionine limitation is caused by degradation of *functional* MetA. This implies that the methionine limitation is brought upon the cell by itself, rather than being caused exclusively by stressors.

If growth arrest protects *E. coli* during heat shocks, and growth arrest is in turn due to methionine limitation, then the addition of methionine should render cells susceptible to heat again. We tested this hypothesis on cultures that were grown, starved, and reinoculated at 44°C in media with and without methionine, and then heat-shocked at 52°C (see ‘Methods’). While cultures without methionine were protected for several hours from heat, cultures with methionine lost more than 99.9% of their viability soon after reinoculation in fresh medium (Fig. 2E).

### Metabolic bistability leads to stable growth arrest

How does methionine limitation trap cells in a state of growth arrest? This is particularly important in order to understand which mechanisms microbes could use to manipulate the fraction of cells entering growth arrest depending on external conditions. While it is clear that degradation of MetA is detrimental for growth, it is important to note how stable growth arrest is: entire cultures can remain growth-arrested for several days, although the conditions would allow for growth in principle (Fig. 1). This stability of growth arrest is puzzling because one would expect that any residual amount of MetA activity, or import of methionine from perished cells, would kickstart growth by supplying methionine, which in turn will be used to synthesize MetA in a self-enhancing positive feedback loop. To answer these questions, we designed a mathematical model of metabolism and protein synthesis that describes the dynamics of internal methionine concentration *c*_met_ and MetA abundance *ϕ*_MetA_. We will use the model to derive a theoretical prediction of lag time, and to perform quantitative tests of our hypotheses.

The model, summarized in Fig. 3A, incorporates processes that affect the cellular methionine level (solid lines) and the MetA protein abundance (dashed lines). Within this model, growth is limited by methionine, Eq. (1), and MetA is degraded at rate *γ*, diluted by growth and synthesized together with biomass at rate *μ*, Eq. (2). Methionine can either be produced by MetA, or taken up from the environment, and is used for protein synthesis and to synthesize S-adenosyl-methionine (SAM), the major methyl donor of the cell, Eq. (3). While much of this SAM is recycled back to methionine in the SAM cycle (Fig. S3), a significant part is used for biosynthesis of polyamines (Table S2), such as spermine and spermidine, which are important for survival in stress conditions ^27–29^. We account for SAM synthesis and other potential losses of methionine flux in a drainage flux (Fig. 3A). In the Supplementary Information we show a detailed derivation of the model and in Table S2 we estimate model parameters based on published measurements.

**Figure 3.**
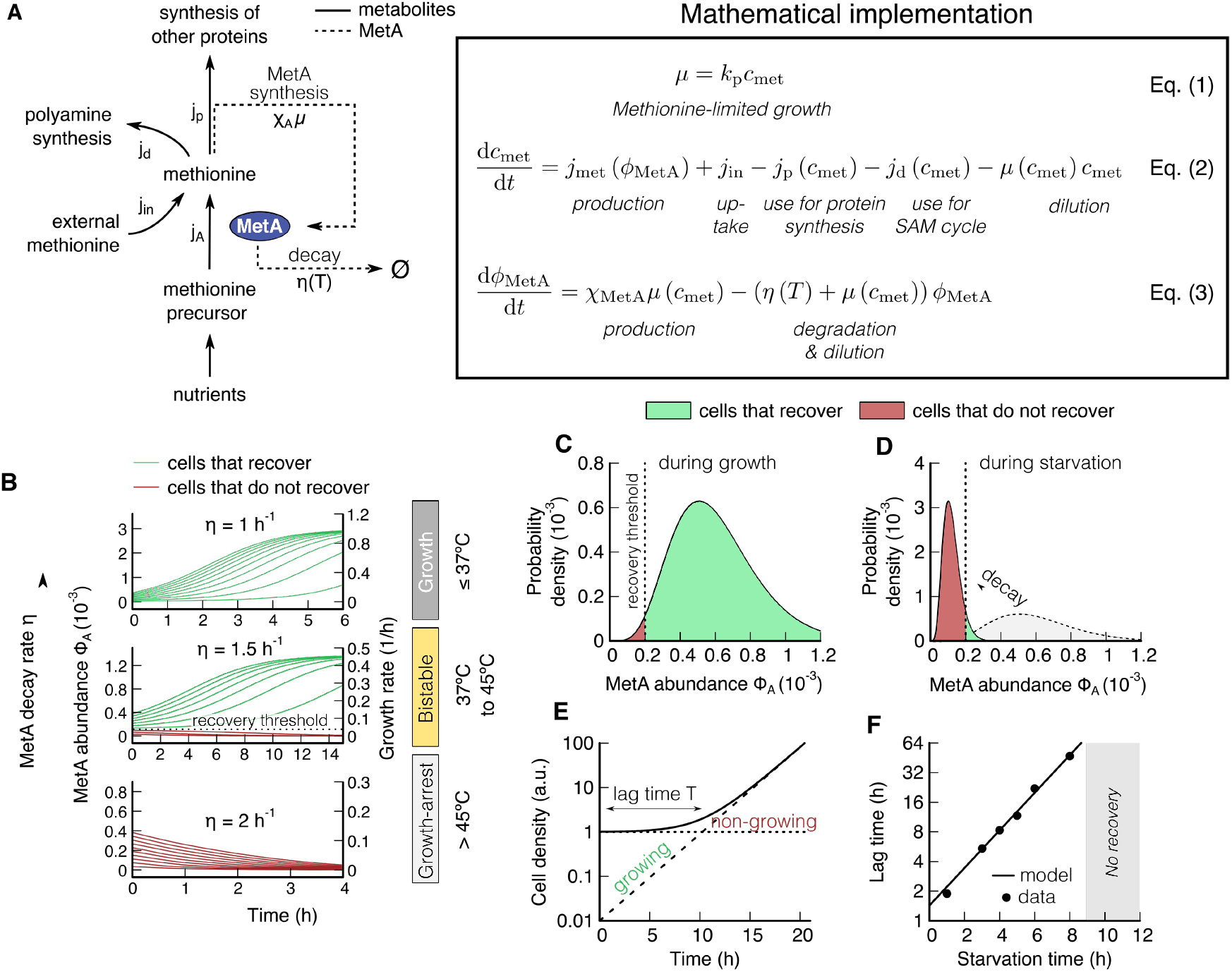
Methionine-MetA model shows that bistability arises from dual use of methionine. *(**A**) Cartoon of the methionine-MetA model. Metabolites: nutrients are taken up, converted first to precursors and finally to methionine by MetA (pink). See Figure S3 for a detailed methionine biosynthesis pathway. External methionine can be imported, if present in the medium. Two fluxes drain methionine, protein synthesis and polyamine synthesis. Proteins: MetA (pink) is synthesized if methionine is present, and will in turn synthesize methionine (positive feedback loop). MetA is degraded at a temperature dependent rate. **(B)** Numerical results of the dynamics of the methionine-MetA model for three different MetA degradation rates (1.0/h, 1.5/h and 2/h). Whether cells can recover depends on the temperature dependent degradation rate. At intermediate temperatures (middle), we observe that only cells with an initial MetA abundance above a recovery threshold can recover (bistable regime). **(C-D)** Distribution of MetA abundance during growth and starvation. Green and red areas indicate cells with MetA abundance that allow (green) and do not allow (red) growth. **(E)** Recovery dynamics of a mixed culture in batch. The time it takes for the growing fraction to overgrow the non-growing fraction is the lag time. (**F**) Theoretical prediction of lag times compared with experiments (45°C). Lag time increases exponentially with starvation time T_lag_* = (1.6 ± 0.3) *exp*(*τ* (0.43 ± 0.02) *h*^−1^). *When cultures are starved for more than 9 hours, they do not recover within three days. The exponential rate fitted, η* = 0.46 *h*^−1^, *is lower than the in-vitro MetA decay rate 1.6 h*^−1^ *measured in Ref. ^27^, possibly due to low levels of ATP that is required for active degradation during starvation ^25^*.

The recovery dynamics of the methionine-MetA model depends on the degradation rate of MetA (Fig. 3B): At low degradation rate, all cells, regardless of their initial state, recover to growth (top panel, green trajectories), while at high degradation rate, none of the cells recover (bottom panel, red trajectories). In between these two extremes, the population splits depending on the initial MetA abundance (Fig. 3B, middle). Cells above a certain ‘recovery threshold’ accumulate MetA and start growing, while cells below the threshold do not. Because both growth and growth-arrest are stable states, this intermediate regime is bistable, and a population can contain cells that are growing and others that are not. Because of the positive-feedback motif between methionine and MetA, the bistability requires a non-linearity in flux^30^, which in our model is caused by the drain of methionine to SAM and disappears if either the drain is shut off (Fig. S4A) or if methionine is supplemented in the medium (Fig. S4B).

Because whether an individual cell recovers or stays dormant depends on its initial MetA abundance, we suspect that fluctuations in MetA abundance cause the split of the population. Protein abundance is indeed highly variable among individual bacteria due to the stochasticity of transcription and translation processes^31,32^, with a typical standard deviation of 40% ^32^. We thus hypothesize that during growth, the vast majority of cells will lie above the recovery threshold (Fig. 3C), and that only due to starvation at elevated temperatures, where MetA will be degraded but not synthesized, the abundance distribution will shift to lower values and many cells will fall below the threshold (Fig. 3D). As a result, only the exponential tail of the MetA abundance distribution will remain above the recovery threshold, leading to a small population of recovering cells. According to the model, the characteristic long lag times observed in Fig. 1 emerge from this mixture of growing and non-growing cells (Fig. 3E).

The model predicts (Supplementary Text 1C) how lag time depends on the fraction of recovering cells, and how degradation of MetA reduces the fraction of recovering cells. We find a simple exponential dependence between lag time *T*_lag_ and the starvation time *τ* and degradation rate *η*,

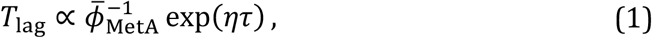

which originates from the exponential tail of MetA abundance, caused by the stochasticity of transcription and translation processes (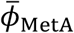 denotes the mean MetA abundance).

A key prediction from the model in Eq. (4) is that the lag time *T*_lag_ should increase exponentially with starvation time *τ*. Because lag time in the model is set by the fraction of the population that recovers, this dependence gives us a direct test of what determines how many cells get trapped in the growth arrest state. Indeed, we can verify the exponential scaling law, and find that it holds over a striking 50-fold range in lag times (Fig. 3F). Cultures starved for more than 9 hours did not recover within 3 days, presumably because all cells of the population are trapped in the growth-arrested state.

The MetA mutants with reduced degradation rates give us the opportunity to test how evolution can tune the fraction of growth-arrested cells, and hence the lag time. The single point mutant I124L, has an experimentally measured degradation rate reduced by 48% compared to wild-type^21^. Using the fit from Fig. 3F, we predict that this mutant should display a lag time of *T*_lag_ = (4.5 ± 1.0) h, which matches the observed (5.2 ± 1.0) h of Fig. 2C well.

### Growth arrest protects cells irrespective of cause

Finally, it is still unclear how heat-induced growth arrest protects cells from subsequent heat-shocks. One possibility is that heat shock proteins, expressed during growth arrest, prepare cells for the upcoming heat shock. To test this hypothesis, we blocked translation during exponential growth by adding chloramphenicol, which leads to growth arrest without new proteins being synthesized ^33^. These experiments were performed at 37°C, where cells are particularly susceptible to subsequent heat shocks. Remarkably, cultures arrested with chloramphenicol survived heat shocks without a significant decrease in viability, whereas the viability of untreated cultures in exponential growth decreased by more than 99.9% (Fig. 4A). Growth arrest by inhibition of protein synthesis thus phenocopies the effect of methionine-limited growth arrest (Fig. 2E). Two alternative types of growth arrest, hyperosmotic shock (3M sorbitol) and sudden carbon starvation (culture washed during exponential growth and resuspended in carbon-free medium, then starved for 2 h, thus preventing adaptation during the stationary phase), also yielded similar protection from heat shocks (Fig. 4A), suggesting that growth arrest by itself, independent of the underlying cause, is protective.

**Figure 4.**
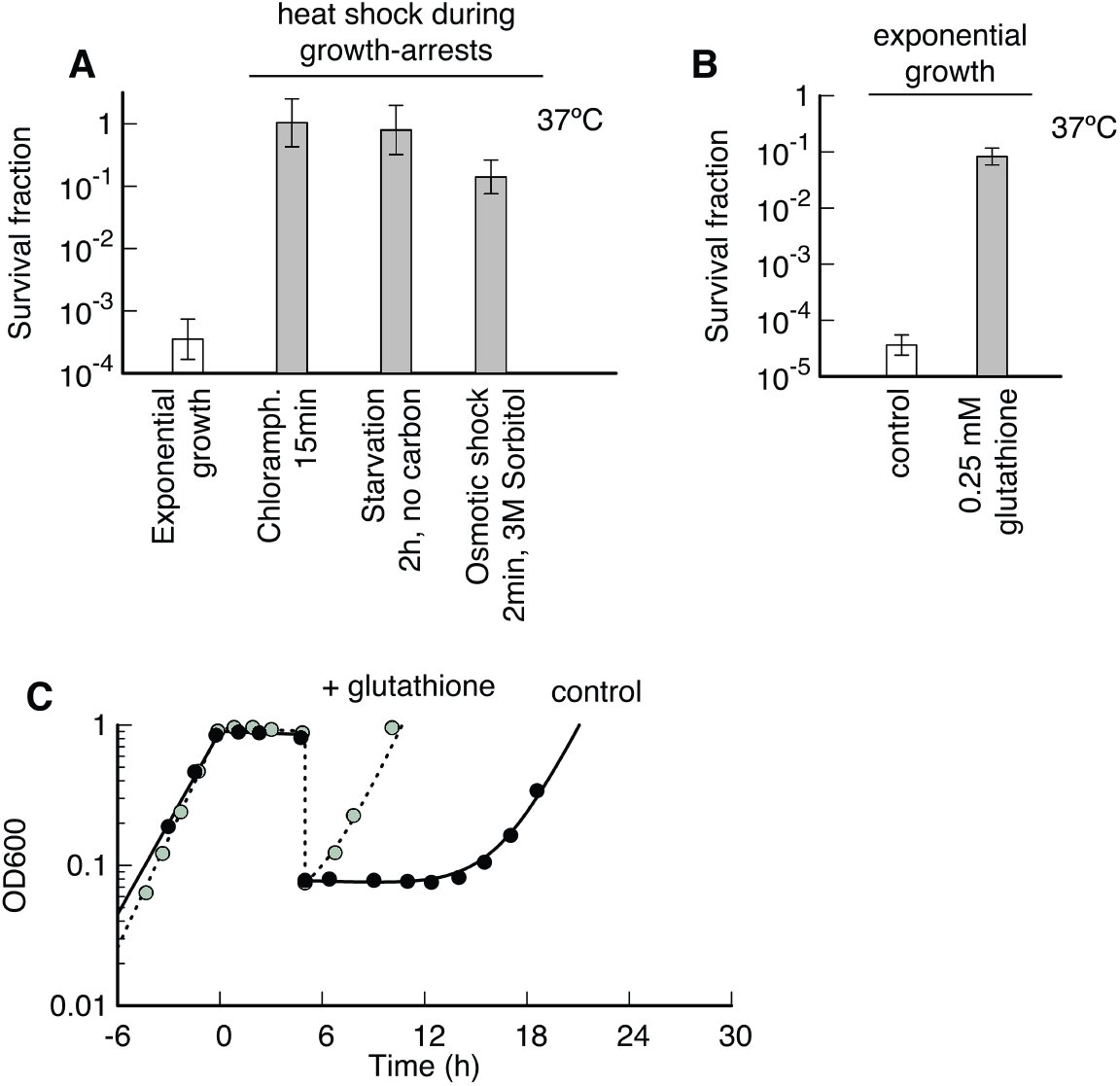
Oxidative stress at elevated temperatures. *(**A**) Killing by heat shock (50°C, 10min) of E. coli cultured at 37°C either in exponential growth (white) or in growth arrest caused by translation inhibition (chloramphenicol), carbon starvation or hyperosmotic shock (grey). Protection from heat killing is independent of the cause of growth arrest. (**B**) Addition of the antioxidant glutathione rescues heat-induced death (50°C, 10 min) in exponential growth indicating that oxidative damage is involved at heat-induced death. (**C**) Addition of glutathione reduces lag times after 5h of starvation from* (*T_lag_* = (1.4 ± 0.2) *h to T_lag_* = (10.3 ± 1.6) *h, N = 3, p = 0.0013)*.

### Oxidative damage contributes to heat-induced death

What are growth-arrested bacteria protected from? For eukaryotic microbes like *Saccharomyces cerevisiae* ^34,35^ and animals like gold fish ^36,37^ it was shown that oxidative damage is involved in heat-induced stress and death. To reduce oxidative damage at high temperatures, *S. cerevisiae* exports antioxidants such as glutathionine ^11^. For bacteria, a similar connection between oxidative stress and temperature stress was hypothesized, based on the observation that temperature sensitive mutants are hypersensitive to oxidative stress ^38^. To test if oxidative stress is indeed involved in heat-induced death, we supplemented cultures in exponential growth at 37°C with glutathione and performed heat shocks. Strikingly, we found that glutathione-supplemented cultures in exponential growth survive 1000-fold better compared to an untreated control (Fig. 4B). This result indicates that growth arrest alleviates lethal oxidative damage that would otherwise kill in heat shocks.

Because oxidative damage is well known to induce aggregation^39,40^ and proteolysis ^41,42^, akin to the aggregation and degradation observed for MetA at high temperatures ^14,15^, we tested if adding the antioxidant glutathione in our standard starvation and reinoculation protocol influences lag times. We found a drastic reduction in lag times for cultures supplemented with glutathione, from 10.4 h to 1.4 h (Fig. 4C). Because lag times are caused by degradation of MetA (Fig. 2), this result suggests that oxidative damage is the trigger for MetA degradation. Similarly, glutathione prevents growth arrest at elevated temperatures in yeast ^11^, suggesting that oxidative damage causes growth arrest more generally in microbes. To test if growth arrest in yeast is also caused by methionine limitation or by a different mechanism, we tested if growth arrest of *Sacharomyces cerevisiae* at 39°C can be rescued with methionine. We found no difference in growth between methionine-supplemented cultures and controls (Fig. S6 A-B), showing that for yeast, growth arrest is caused by a different mechanism. The protective feature of growth arrest, however, is shared by yeast: While growing yeast (30°C) died by a factor of 500 during heat shocks, starving yeast largly survived (Fig S6 C). Taken together, these findings show that employing growth arrest to combat heat-induced death may be a more general strategy of microbes.

## DISCUSSION

We have shown that breakdown of methionine biosynthesis at elevated temperatures causes growth arrest in *Escherichia coli*. Despite its fitness cost under habitable conditions, growth arrest is beneficial for *E. coli* because it protects cells from heat-induced death when temperatures rise beyond 50°C. We demonstrated that this growth arrest can be alleviated by simple point mutations and knock-outs. We therefore suggest that it constitutes a self-controlled growth arrest that has likely evolved as an active response to heat stress, orthogonal to the well-studied heat shock response.

### Relation to ‘viable but not culturable’ (VBNC) and persistence states

Growth arrest similar to the phenomenon described in our work is frequently encountered in a variety of microbes across the phylogenetic spectrum ^43^ when exposed to stress conditions outside of the optimal culture conditions ^44,45,43^. A growing body of literature suggests the importance of growth arrest in the context of ‘viable but not culturable’ (VBNC) states^43,45^ and persistence^24,46^, in particular in food safety^47,48^ and antibiotic treatment^24^. However, pinning down mechanisms of how bacteria achieve growth arrest and why such mechanisms have evolved has proven challenging ^49,50^. Our results show that proteolytic regulation of basic biosynthetic pathways is an efficient strategy to robustly induce growth-arrest.

### Shut-off of methionine synthesis is ideally suited to halt growth

While a lack of any amino acid would stop translation, lack of methionine specifically halts translation initiation, because the majority of bacterial start codons code for N-formylmethionine (fMet), a direct derivative of methionine. Lack of other amino acids would halt elongation after initiation, leading to protein fragments that would need to be recycled by the cell. Furthermore, inhibition of translation triggers the stringent response alarmone (p)ppGpp ^51^. This global regulator in bacteria inhibits synthesis of rRNA and tRNA ^52^, transcription ^53^, as well as DNA replication ^54,55^, leading to a global shut-off in macromolecule synthesis. At the same time, the central metabolism and generation of energy-rich molecules are upstream of MetA, so that cells are presumably well fueled to perform energy-dependent maintenance. This combination of inhibited biosynthesis, but non-inhibited metabolism makes methionine an ideal bottleneck to induce growth-arrest in bacteria, which lack global regulators of growth, and we therefore speculate that this is the reason the for conservation of temperature-dependent methionine synthesis breakdown in bacteria across the phylogenetic spectrum ^17,19,20,18^.

### MetA is a biological ‘thermal fuse’

The temperature-induced degradation of MetA has a counterpart in electrical engineering: electrical devices are protected from excessive heat with ‘thermal fuses’, which cut off electric current by physically disintegrating, typically by melting. The fuse has the role of an emergency protection, kicking in when other regulatory mechanisms have failed, in order to protect expensive components. In this analogy, first suggested by Price-Carter et al ^56^ for acid stress, we interpret MetA as a biological thermal fuse. If the benefit of increasing survival chances outweighs the cost of increased synthesis and missed potential for growth, then the thermal fuse will have a net positive effect on fitness. The high expression levels combined with high degradation rates, as in the case of MetA, reflects a highly volatile strategy, typical for biological regulators ^22^. In fact, it is ideally suited to stressful conditions to ensure fast growth if conditions are bearable, while also quickly shutting off biosynthesis if conditions worsen.

### Bet-hedging between growth and survival in anticipation of rare catastrophic events

The bistability in growth arrest could play a vital role in optimizing fitness at elevated temperatures. Because both growth and growth arrest are stable, the bacterial population can split into two subpopulations. If individual cells can switch between states of growth and arrest depending on the external environment and their internal methionine concentration, then according to the model, the fitness of the population is optimized by partitioning the population into two sub-populations; a growing and growth-arrested population. Such a mixture of two distinct behaviors in a genetically homogeneous population of cells is a hallmark of bet-hedging strategies, typical for biolgical systems in unpredictable environments ^57–59^. Bet-hedging aims to increase long-term fitness beyond what would be possible for either pure strategies of growth and growth arrest. Thus having a growth-arrested subpopulation will allow microbes to survive sudden heat shocks, while at the same time proliferating at sub-critical temperatures – enabling them to robustly thrive at the edge of habitable conditions.

## Acknowledgments

This work was supported by the German Research Foundation (DFG) via the Excellence Cluster ORIGINS (through U.G.), and by the Life initiative of the Volkswagen Foundation (through U.G.). E.B. was supported by a DFG fellowship through the Graduate School of Quantitative Biosciences Munich (QBM). S.J.S was supported by a Long-Term Fellowship (ALTF 782-2017) from the European Molecular Biology Organization (EMBO) and Long-Term Fellowship (LT000597/2018-L) from the Human Frontiers in Science Program (HFSP).

## Author contributions

SJS and UG conceived this study. SJS, EB, MGH and ZG designed and performed experiments. YFC constructed strains. SJS, MB and UG developd the theoretical model. SJS, EB, MB and UG wrote the paper. MB and UG funded and supervised the project.

## Data availability statement

Source data of figures are available upon request from the corresponding authors.

## Code availability

The numerical implementation of the theoretical model used to generate Figure 3 is available upon request from the coresponding authors.

## METHODS

### Contact for Reagent and Resource Sharing

Further information and requests for resources and reagents should be directed to and will be fulfilled by the lead contact.

### Strains

All strains used in this study are derived from wild type *E. coli* K-12 strain NCM3722 ^60^. Amino acid substitutions MetA_I229Y_ and MetA_LYD_ (containing point mutations I124L-I229Y-N267D) in strains I229Y and LYD ^21^ were transferred to NCM3722 ΔMetA via P1 transduction to yield strains YCE55 (NCM3722 metA_I229Y_) and YCE57 (NCM3722 metA_LYD_), using methionine as a selective marker.

### Culture media

The culture medium for *E. coli* used in this study is based on N^−^C^−^ minimal medium ^61^, containing K_2_SO_4_ (1 g), K_2_HPO_4_·3H_2_O (17.7 g), KH_2_PO_4_ (4.7 g), MgSO_4_·7H_2_O (0.1 g) and NaCl (2.5 g) per liter. The medium was supplemented with 20 mM NH_4_Cl, as nitrogen source, and 5 to 10 mM glycerol, as the sole carbon source. *S. cerevisiae* were cultured in yeast nitrogen base medium (without amino acids) and supplemented with 0.5% glucose. All chemicals were purchased from Carl Roth, Karlsruhe, Germany. When needed, 0.1 mg/ml ampicillin (stock prepared fresh and stored at −20°C) and/or 0.067 mM methionine (stock stored at 4°C) were added to the culture

### Growth protocol for E. coli

Cells were taken from a −80°C glycerol stock, streaked out on a LB agar plate and incubated around 12 hours at 37°C. A single colony was picked and grown in batch culture. For temperatures up to 39°C, cells were grown first on LB at the desired temperature (seed culture). Before reaching starvation, they were diluted and inoculated into pre-warmed minimal medium supplemented with glycerol (pre-culture). Cells grew in the pre-culture for several doublings to ensure exponential growth and were then diluted and inoculated into pre-warmed glycerol minimal medium (experimental culture). Inoculation into the latter was chosen such that cells perform at least three additional doublings in the experimental culture before growth was measured. For high temperatures (above 39°C), the seed culture medium used was glycerol minimal medium with 1% LB in order to help the cells transition into the pre-culture.

For small culture volumes, 5 ml of culture were grown in 20 mm × 150 mm glass test tubes (Fisher Scientific, Hampton, NH, USA) with disposable, polypropylene Kim-Kap closures (Kimble Chase, Vineland, NJ, USA). Larger volumes, 25 ml, 50 ml and 100 ml were grown in 125 ml, 250 ml and 500 ml baffled Erlenmeyer flasks (Carl Roth, Karlsruhe, Germany), respectively, and Kim-Kap closures. All cultures were grown in a shaking water bath (WSB-30, Witeg, Wertheim, Germany) with water bath preservative (Akasolv, Akadia, Mannheim, Germany).

### Growth protocol for S. cerevisiae

Cells were streaked from a −80°C glycerol stock onto a YPD agar plate and incubated for approximately 30 hours at 30°C. A single colony was picked and grown in batch culture with 4 mL NBM containing 1% YPD. Where necessary, the culture was supplemented with methionine (0.067 mM). All cultures were incubated in a shaking water bath with the culture tubes angled parallel to the direction of shaking. The culture tubes and shaking water bath used were identical to those used for the cultivation of *E. coli*.

### Protocol to induce long lag times at high temperatures

Bacteria were first grown in minimal medium supplemented with glycerol as a carbon substrate, as described above, at the desired temperature (37 to 45°C). After carbon depletion, cultures starved for 1 h to 24 h. Then, cultures were diluted into fresh N^−^C^−^ supplemented with glycerol and NH_4_Cl and allowed to recover.

### Quantification of lag time

The lag time after glycerol re-addition was extracted from optical density measurements, and corrected for the loss of viability during the period of starvation. Growth was modeled as OD_600_ = *A* + *Be^μt^*, which for long times converges to 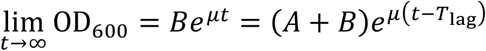, with lag time *T*_lag_. We extracted growth rate *μ* and the growing fraction of cells, *B*, from the time derivative of optical density d OD_600_/d*t* = *μBe^μt^*, using a least square fit across the recovery regime where the time derivative increases exponentially. We measure (*A* + *B*) as the optical density after dilution. The lag time *T*_lag_ = −*μ*^−1^ ln(*B*/(*A* + *B*)), can then be calculated from the extracted parameters. Lastly, because optical density measurements do not distinguish between living and dead cells in the culture, and a fraction of the culture has perished during starvation, we corrected the lag time estimation by subtracting the number of dead cells *N*_d_ from the initial number *N*(0) = *A* + *B*. This gives a correction factor 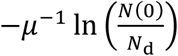. For 5 h starvation at 45°C, about one third of the cells perish. Thus, with growth rate 0.45/h at 45°C, this yields a correction factor of 0.9 h. In the main text, all lag times are corrected for perished cells. Note that the bi-exponential fits in Fig. 1, that include dynamics of both death and growth, are not used to extract lag times.

### Time-lapse microscopy

Microscopy was performed using agar pads in glass-bottom dishes. Agar pads contained medium identical to the culture medium, N^−^C^−^ supplemented with 20 mM NH_4_Cl, 20 mM Glycerol and 2 % low melting point agarose (Carl Roth, Karlsruhe, Germany). Cells were diluted and pipetted on the prewarmed agar pad and kept in the incubator. After the surface of the agar pad had dried, the agar pad was flipped on the glass window of a prewarmed glass bottom dish. The glass bottom dishes were sealed with parafilm to reduce evaporation.

Time-lapse microscopy was performed using a Nikon Ti-E microscope controlled by NIS-Elements (Nikon, Tokyo, Japan). We used a phase contrast Plan Apo 100x oil objective, with numerical aperture of 1.45 and a refractive index of 1.515 (Nikon, Tokyo, Japan) and a SCMOS camera (Zyla-5.5, Andor, Belfast, Northern Ireland) with a binning of 1×1, a readout rate of 200 MHz and an exposure time of 200 ms. Conversion gain was set on 1/3 Dual gain and the spurious noise filter was activated. The calibration from length units to pixels was defined as 0.07 μm/px. Measurements were performed with activated perfect focus system and a PriorScan III drive stage. Temperature control was set by the Okolab Cage Incubation System and confirmed with manual temperature measurements. Cell growth was monitored for 24 hours with continuous acquisition of phase-contrast and fluorescence images every 30 min at the desired temperature, e.g. 45°C.

### Live/dead stain

Commercial BacLight®LIVE/DEAD (Thermo Fisher Scientific Inc., Waltham, Massachusetts, USA) staining was used when cells were microscopically imaged, according to manufacturing specifications.

### Heat shock protocol

Heat shocks were performed by transferring batch cultures (5 ml culture volume in a 20 mm x 150 mm borosilicate glass tube (Fisher Scientific, NH, USA)) from one water bath to another (WSB-30, Witeg, Wertheim, Germany). Water baths were pre-set and equilibrated at the desired temperatures.

### Survival quantification

Survival in starvation was quantified by counting colony forming units (CFU). Samples were diluted in carbon free N^−^C^−^ minimal medium and spread on LB agar plates using Rattler Plating Beads (Zymo Research, Irvine, CA, USA) with a target density of 200 colonies per plate. LB agar was supplemented with 25 μg/ml 2,3,5-triphenyltetrazolium chloride to stain colonies and increase contrast for automated colony counting (Scan 1200, Interscience, Saint-Nom-la-Bretèche, France). Staining or automation of counting had no significant effect on viability measurements or accuracy, compared to unstained, manually counted samples (<1% systematic error). Petri dishes were 92 × 16 mm (Sarstedt, Nümbrecht, Germany) and agar plates were incubated for 12 hours at 37°C to ensure optimal colony size.

### Numerical analysis of the model

The numerical solution in Fig. 3B was obtained by numerically integrating Eqs. (1-3) using the nonstiff differential equation solver *ode45* in Matlab (Mathworks, Natick, MA, USA). Initial conditions were *c*_met_(0) = 0 and *ϕ*_MetA_(0) ranging from 2 · 10^−5^ to 2 · 10^−4^. The parameters used in the calculations were *a*_1_ = 4.06 M^−1^, *a*_2_ = 3.1 · 10^−5^, *a*_3_ = 148 M^−2^, *χ*_MetA_ = 2.79 · 10^−3^, *K*_M_ = 10^−6^ M, see Table S2.

The phase diagrams in Fig. S4 were calculated by solving Eq. (S12) using the *solve* function in Matlab (Mathworks, Natick, MA, USA). We shaded the regions according to the number of stable fixed-points at concentrations higher or equal to zero. One solution at 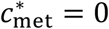 (stable) was denoted ‘non-growing region’. Two zeros, 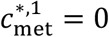 (unstable) and 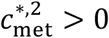 (stable) was denoted ‘growth region’. Three zeros was denoted as the ‘bistable region’. If methionine influx was included, Fig. S4E, one stable solution at 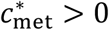 means that cells can grow on external methionine. This region was shaded ‘grey to white’ to symbolize the growth rate dependence on the methionine influx. The parameters used in the calculations were *a*_1_ = 4.06 M^−1^, *a*_2_ = 3.1 · 10^−5^, *a*_3_ = 148 M^−2^, *χ*_MetA_ = 2.79 · 10^−3^, *K*_M_ = 10^−6^ M, see Table S2.

### Statistical analysis

Lag times with individual tests, e.g. methionine versus no methionine, were performed in 2-3 biological repeats as mentioned in figure captions and are reported with one standard deviation. Measurements that are quantitative tests, e.g. influence of starvation time, were performed as single repeats, at multiple different perturbation strength. Results of fits are reported with standard error. Measurements of cell survival to antibiotic treatment were performed in duplicates for each temperature. Survival measurements upon temperature shift from 37, 40, 42 and 45 °C to 50°C were performed at least twice for wild-type and MetA stabilized mutants.

