## Supplemental Information for "MetA is a ‘thermal fuse’ that arrests growth and protects *Escherichia coli* at elevated temperatures"

[3] Contributed equally

### *I. Supplementary derivations*

In this section of the supplementary information, we will derive theoretical solutions for the model discussed in the main section. In particular, we focus here on the stability analysis, the MetA recovery threshold and the derivation of the lag times. In addition, we make a detailed estimation of the theoretical parameters from measurements in the literature.

#### *A. Stability analysis*

In this section we discuss the stability of the non-linear equation system proposed in Eqs. (1-3) of the main text. The standard approach to stability analysis requires derivation of the two nullclines of the system, where 1) the methionine concentration and 2) the active MetA abundance are constant [1]. At the intersection of the two nullclines, we find the 'fixed points'. When the system is evolving, the cells will relax towards the stable fixed points (and away from the unstable fixed points). In our case, the possible stable fixed-point positions are growth or growth arrest and their existence will be discussed in the section thereafter. We will first derive the nullclines and then the criteria for the existence of the fixed points.

##### *1. Derivation of the nullclines*

In Eq. (2) of the main text we derived the dynamics of the internal methionine concentration as

$$\frac{\mathrm{d}c_{\mathrm{met}}}{\mathrm{d}t} = j_{\mathrm{MetA}}(\phi_{\mathrm{MetA}}) + j_{\mathrm{in}} - j_{\mathrm{p}}(c_{\mathrm{met}}) - j_{\mathrm{d}}(c_{\mathrm{met}}) - \mu c_{\mathrm{met}}, \tag{S1}$$

With the individual contributions being

$$j_{\mathrm{MetA}} = h\phi_{\mathrm{MetA}} \tag{S2}$$

$$j_{\mathrm{p}} = c_{\mathrm{met}}^{\mathrm{pro}} k_{\mathrm{p}} c_{\mathrm{met}} = c_{\mathrm{met}}^{\mathrm{pro}} \mu \tag{S3}$$

$$j_{\mathrm{d}} = k_{\mathrm{d}}\, u(c_{\mathrm{met}}) \tag{S4}$$

$$j_{\mathrm{in}} = \mathrm{const} > 0 \tag{S5}$$

and

$$\mu = k_\mathrm{p} c_\mathrm{met}, \quad \text{(S6)}$$

$$u(c_\mathrm{met}) = c_\mathrm{met}/(K_\mathrm{M} + c_\mathrm{met}). \quad \text{(S7)}$$

The production flux $j_\mathrm{MetA} = h\phi_\mathrm{MetA}$ equals the production rate $h$ times the MetA abundance $\phi_\mathrm{MetA}$, because MetA is the limiting step in synthesis[2]. Here, MetA abundance $\phi_\mathrm{MetA}$ is defined as the fraction of the proteome that is MetA. Uptake flux $j_\mathrm{in}$ depends on the presence of methionine and metabolites containing methionine in the medium. The methionine flux to protein synthesis $j_\mathrm{p} = c_\mathrm{met}^\mathrm{pro}\,\mu$ is calculated by assuming that cells maintain a constant protein density, in which case the flux needs to be sufficient to prevent dilution of the 'methionine concentration bound in proteins' $c_\mathrm{met}^\mathrm{pro}$, if the cell is growing at rate $\mu$. We estimate an internal methionine concentration of $c_\mathrm{met}^\mathrm{pro} = 27.5$ mM from the codon frequency of methionine (3.2%) and a typical protein concentration of about 0.125 g/ml, see Table S2.

Finally, we consider that methionine is consumed for S-adenosyl-methionine (SAM) synthesis, the major methyl donor of the cell. See Fig. S3 for a summary of the methionine pathway. While much of this SAM is recycled back to methionine, a part of it is used for biosynthesis of polyamines, such as spermine and spermidine, which are important for survival in stress conditions [3–5]. Of these metabolites, spermidine is present at a particularly high intracellular concentration, $c_\mathrm{spe} = 1.4$ mM [6]. Because one methionine is consumed per spermidine, we can estimate the fraction of the methionine flux that is used for spermidine synthesis as $c_\mathrm{spe}/c_\mathrm{met}^\mathrm{pro} = 1.4\ \mathrm{mM}/27.5\ \mathrm{mM} = 5\%$. Since polyamine synthesis is an overall small part of the cell's biomass, we assume that the 'drainage flux', $j_\mathrm{d}$, saturates at some level and model it as a saturating function of the internal methionine concentration, $j_\mathrm{d} = k_\mathrm{d}\ c_\mathrm{met}/(c_\mathrm{met} + K_\mathrm{M})$, with $k_\mathrm{d}$ being the maximum efflux rate and $K_\mathrm{M}$ the Michaelis constant. Efflux of methionine from the cell, either by passive diffusive loss or by active export, could also contribute to the drain of internal methionine, but will be neglected here. If existent, they would effectively increase the drainage flux $j_\mathrm{d}$. The combination of all metabolite fluxes determines the dynamics of the methionine concentration, the limiting bottleneck at elevated temperatures.

The nullcline of the methionine concentration is defined as the manifold in the space of $\phi_\mathrm{MetA}$ and $c_\mathrm{met}$, in which $\mathrm{d}c_\mathrm{met}/\mathrm{d}t = 0$. Setting Eq. (S1) to zero, we get

$$\frac{\mathrm{d}c_{\mathrm{met}}}{\mathrm{d}t} = f_1(\phi_{\mathrm{MetA}}, c_{\mathrm{met}}) = h\phi_{\mathrm{MetA}} + j_{\mathrm{in}} - c_{\mathrm{met}}^{\mathrm{pro}} k_{\mathrm{p}} c_{\mathrm{met}} - k_{\mathrm{d}}\, u(c_{\mathrm{met}}) - k_{\mathrm{p}} c_{\mathrm{met}}^2 = 0. \quad \text{(S8)}$$

By solving this equation for $\phi_{\mathrm{MetA}}$ and defining new parameters $a_1 = c_{\mathrm{met}}^{\mathrm{pro}} k_{\mathrm{p}} h^{-1}$, $a_2 = k_{\mathrm{d}} h^{-1}$ and $a_3 = k_{\mathrm{p}} h^{-1}$, as well as $\tilde{j}_{\mathrm{in}} = j_{\mathrm{in}} h^{-1}$, we derive the nullcline of the methionine concentration,

$$g_{\mathrm{met}}(c_{\mathrm{met}}) = \phi_{\mathrm{MetA}} = a_1 c_{\mathrm{met}} + a_2 u(c_{\mathrm{met}}) + a_3 c_{\mathrm{met}}^2 - \tilde{j}_{\mathrm{in}}. \quad \text{(S9)}$$

The second component of the model is the dynamics of the MetA abundance, $\phi_{\mathrm{MetA}}$. MetA is synthesized during growth at rate $\chi_{\mathrm{MetA}}\mu$, where $\chi_{\mathrm{MetA}}$ is the fraction of the newly synthesized proteome that is MetA. Since we are interested in the regime of low methionine abundance, we assume that $\chi_{\mathrm{MetA}}$ is set constantly to full expression. In addition to dilution by growth, $\mu\phi_{\mathrm{MetA}}$, the MetA abundance decays at a constant rate, $\eta(T)\phi_{\mathrm{MetA}}$ [7,8]. The decay rate $\eta(T)$ is temperature $T$ dependent [8], ATP dependent [7] and includes any process that renders MetA unfunctional, from degradation [7] to thermal aggregation [9]. Combined, the dynamics of the MetA abundance gives Eq. (3) from the main text, which we set to zero to obtain the nullcline

$$\frac{\mathrm{d}\phi_{\mathrm{MetA}}}{\mathrm{d}t} = f_2(\phi_{\mathrm{MetA}}, c_{\mathrm{met}}) = \chi_{\mathrm{MetA}}\mu(c_{\mathrm{met}}) - \big(\eta(T) + \mu(c_{\mathrm{met}})\big)\phi_{\mathrm{MetA}} = 0. \quad \text{(S10)}$$

By solving for $\phi_{\mathrm{MetA}}$, we obtain the nullcine for the active MetA abundance to be

$$g_{\mathrm{MetA}}(c_{\mathrm{met}}) = \phi_{\mathrm{MetA}} = \chi_{\mathrm{MetA}} \frac{c_{\mathrm{met}}}{\tilde{\eta} + c_{\mathrm{met}}}, \quad \text{(S11)}$$

where we define a new variable $\tilde{\eta} = \eta k_{\mathrm{p}}^{-1}$ and use $\mu = k_{\mathrm{p}} c_{\mathrm{met}}$ from Eq. (1) of the main text.

#### 2. *Stability criteria*

The fixed points of the system are found at the intersection of both nullclines $g_{\mathrm{met}}(c_{\mathrm{met}}) = g_{\mathrm{MetA}}(c_{\mathrm{met}})$, i.e.

$$\chi_{\mathrm{MetA}} \frac{c_{\mathrm{met}}^{*}}{\tilde{\eta} + c_{\mathrm{met}}^{*}} = \phi_{\mathrm{MetA}} = a_1 c_{\mathrm{met}}^{*} + a_2 u(c_{\mathrm{met}}^{*}) + a_3 c_{\mathrm{met}}^{*2} - \tilde{j}_{\mathrm{in}}, \tag{S12}$$

where $c_{\mathrm{met}}^{*}$ is the fixed-point of the internal methionine concentration. The fixed-point of the MetA abundance $\phi_{\mathrm{MetA}}^{*}$ can then be obtained by plugging in $c_{\mathrm{met}}^{*}$ into either nullcline. If we then solve the fixed-points of Eq. (S12) numerically, we can get an exact solution for the growth fixed point. This numerical solution is shown in the main text in Figures 5D, E.

Depending on the parameter regime, there are either one, two or three fixed-points in the regime $c_{\mathrm{met}} \geq 0$ and $\phi_{\mathrm{MetA}} \geq 0$. We can get vital information about the system by calculating stability criteria analytically. We consider the simplest case of no external methionine influx $\tilde{j}_{\mathrm{in}} = 0$, where the trivial solution of the fixed-point equation, Eq. (S12), is $c_{\mathrm{met}}^{\mathrm{d}} = 0$ and $\phi_{\mathrm{MetA}}^{\mathrm{d}} = 0$. This fixed-point corresponds to dormancy, thus the superscript 'd', because in this case the steady state growth rate is zero, $\mu^{\mathrm{d}} = k_{\mathrm{p}} c_{\mathrm{met}}^{\mathrm{d}} = 0$. This fixed-point can either be stable, i.e. the cell will stop growing when close to this fixed-point, or unstable, i.e. the cell will start growing, even at very low $c_{\mathrm{met}}$ and $\phi_{\mathrm{MetA}}$ levels. The stability can be checked using the Jacobian matrix of the system, defined as

$$J = \begin{pmatrix} \frac{\partial f_1}{\partial c_{\mathrm{met}}} & \frac{\partial f_1}{\partial \phi_{\mathrm{MetA}}} \\ \frac{\partial f_2}{\partial c_{\mathrm{met}}} & \frac{\partial f_2}{\partial \phi_{\mathrm{MetA}}} \end{pmatrix} \tag{S13}$$

At the trivial fixed point $c_{\mathrm{met}}^{\mathrm{d}} = 0$ and $\phi_{\mathrm{MetA}}^{\mathrm{d}} = 0$, we find

$$J = \begin{pmatrix} -h(a_1 + a_2/K_{\mathrm{M}}) & h \\ \chi_{\mathrm{MetA}} k_{\mathrm{p}} & -\eta \end{pmatrix}. \tag{S14}$$

The stability of the fixed point can then be checked by computing the trace

$$\tau = \frac{\partial f_1}{\partial c_{\mathrm{met}}} + \frac{\partial f_2}{\partial \phi_{\mathrm{MetA}}} \tag{S15}$$

and determinant

$$\Delta = \frac{\partial f_1}{\partial c_{\mathrm{met}}} \frac{\partial f_2}{\partial \phi_{\mathrm{MetA}}} - \frac{\partial f_1}{\partial \phi_{\mathrm{MetA}}} \frac{\partial f_2}{\partial c_{\mathrm{met}}} \tag{S16}$$

of the Jacobian matrix [1]. Because the trace $\tau = -(h(a_1 + a_2/K_{\mathrm{M}}) + \eta)$ is negative, the stability is determined solely by the sign of the determinant $\Delta = h(a_1 + a_2/K_{\mathrm{M}})\eta - h\chi_{\mathrm{MetA}}k_{\mathrm{p}}$. If $\Delta > 0$, then the dormancy fixed-point is stable. This condition is met if

$$\eta > \frac{\chi_{\mathrm{MetA}} k_{\mathrm{p}}}{(a_1 + a_2/K_{\mathrm{M}})}, \tag{S17}$$

which can be formulated in terms of the original parameters as

$$\eta > \frac{\chi_{\mathrm{MetA}} h}{c_{\mathrm{met}}^{\mathrm{pro}} + k_{\mathrm{d}}/(k_{\mathrm{p}} K_{\mathrm{M}})}. \tag{S18}$$

To get an analytical estimate for the stability criterium of the growth fixed-point, we make several simplifying assumptions. We first note that the MetA abundance nullcline $g_{\mathrm{MetA}}(c_{\mathrm{met}})$ levels to a constant, $g_{\mathrm{MetA}}(c_{\mathrm{met}} \to \infty) \to \chi_{\mathrm{MetA}} = \mathrm{const}$, for high methionine concentrations, see Eq. (S10), while the methionine nullcline $g_{\mathrm{met}}(c_{\mathrm{met}})$ increases monotonically to infinity in the same limit, $g_{\mathrm{met}}(c_{\mathrm{met}} \to \infty) \to \infty$, see Eq. (S8). Thus, if at low methionine concentrations, the methionine nullcline $g_{\mathrm{met}}(c_{\mathrm{met}})$ is lower than the MetA abundance nullcline $g_{\mathrm{MetA}}(c_{\mathrm{met}})$, these two nullclines will cross and create a fixed-point at finite growth, $c_{\mathrm{met}}^{\mathrm{g}} > 0$ and $\phi_{\mathrm{MetA}}^{\mathrm{g}} > 0$, where the superscript ‘g’ stands for growth.

There are two possibilities that the methionine nullcline $g_{\mathrm{met}}(c_{\mathrm{met}})$ can be lower than the MetA abundance nullcline $g_{\mathrm{MetA}}(c_{\mathrm{met}})$. The first option is that for very low methionine concentration, $0 < c_{\mathrm{met}} \ll K_{\mathrm{M}}$, the methionine nullcline increases less than the MetA abundance nullcline, $\mathrm{d}g_{\mathrm{met}}/\mathrm{d}c_{\mathrm{met}} < \mathrm{d}g_{\mathrm{MetA}}/\mathrm{d}c_{\mathrm{met}}$. Testing this was the essence of the stability analysis done for the non-growing fixed-point in this section: if the dormancy fixed point is stable, $\mathrm{d}g_{\mathrm{met}}/\mathrm{d}c_{\mathrm{met}} >$

$\mathrm{d}g_{\mathrm{MetA}}/\mathrm{d}c_{\mathrm{met}}$; if the dormancy fixed point is unstable, $\mathrm{d}g_{\mathrm{met}}/\mathrm{d}c_{\mathrm{met}} < \mathrm{d}g_{\mathrm{MetA}}/\mathrm{d}c_{\mathrm{met}}$. The second option is that, at a higher methionine concentration, the methionine nullcline $g_{\mathrm{met}}(c_{\mathrm{met}})$ is again lower than the MetA nullcline $g_{\mathrm{MetA}}(c_{\mathrm{met}})$. We estimate the regime where this happens, by comparing the two nullclines at relatively low methionine concentrations $K_{\mathrm{M}} \ll c_{\mathrm{met}} \ll c_{\mathrm{met}}^{\mathrm{g}}$, in the regime where the drain has already saturated, i.e., $u(K_{\mathrm{M}} \ll c_{\mathrm{met}}) = 1$, but the methionine concentration is still low compared to the steady state, $c_{\mathrm{met}}^{\mathrm{g}}$.

In this regime, the methionine nullcline can be approximated as $g_{\mathrm{met}}(c_{\mathrm{met}}) \approx a_1 c_{\mathrm{met}} + a_2$, by omitting quadratic terms in Eq. (S8). The MetA abundance nullcline in the same regime can be approximated as, $g_{\mathrm{MetA}}(c_{\mathrm{met}}) \approx \chi_{\mathrm{MetA}} \tilde{\eta}^{-1}\, c_{\mathrm{met}}$, when we omit higher order terms in Eq. (S10). Combined, these two approximations yield that, if $a_1 c_{\mathrm{met}} + a_2 < \chi_{\mathrm{MetA}} \tilde{\eta}^{-1}\, c_{\mathrm{met}}$, then the methionine nullcline lies below the MetA nullcline.

If we take $a_2 \approx 0$, then we can estimate an upper limit,

$$\tilde{\eta} < \frac{\chi_{\mathrm{MetA}}}{a_1} \tag{S19}$$

or in the original parameters,

$$\eta < \frac{\chi_{\mathrm{MetA}} h}{c_{\mathrm{met}}^{\mathrm{pro}}}, \tag{S20}$$

for the degradation rate in this regime.

From the above analysis we conclude that three distinct regimes exist. First, if

$$\eta < \frac{\chi_{\mathrm{MetA}} h}{c_{\mathrm{met}}^{\mathrm{pro}} + k_{\mathrm{d}}/(k_{\mathrm{p}} K_{\mathrm{M}})}, \tag{S21}$$

then only the growing state is stable, and all cells in the population will grow. Second, if

$$\frac{\chi_{\mathrm{MetA}} h}{c_{\mathrm{met}}^{\mathrm{pro}} + k_{\mathrm{d}}/(k_{\mathrm{p}} K_{\mathrm{M}})} < \eta < \frac{\chi_{\mathrm{MetA}} h}{c_{\mathrm{met}}^{\mathrm{pro}}} \tag{S22}$$

then both a stable growth and dormancy fixed-points exists, and the culture can split into two subpopulations. Third, if

$$\eta > \frac{\chi_{\mathrm{MetA}} h}{c_{\mathrm{met}}^{\mathrm{pro}}} \tag{S23}$$

then only the dormant state is stable and the entire population will converge to growth arrest.

##### *B. Calculation of the recovery threshold*

In the case that the environment permits two stable fixed-points, growth and dormancy, the question is: in which state will an individual cell end? To address this question, we will calculate the necessary MetA abundance for recovery, called the "recovery threshold" in the main text. Above this threshold, the cell will have enough MetA to start producing more methionine and, conversely, more MetA. Below the threshold, too much methionine is drained, the degradation of MetA is faster than the production of new MetA and the cells will end in a stable state of dormancy.

###### *1. Separation of time-scales*

In starvation, according to the model, the internal metabolite concentrations will drop quickly, i.e. $c_{\mathrm{met}} \approx 0$, and the MetA abundance will decay slowly,

$$\left.\frac{\mathrm{d}\phi_{\mathrm{MetA}}}{\mathrm{d}t}\right|_{c_{\mathrm{met}}=0} = -\eta\phi_{\mathrm{MetA}}. \tag{S24}$$

During this decay, at some point the cell will cross a threshold $\phi_{\mathrm{MetA}}^{\mathrm{th}}$, below which it cannot recover.

To estimate the dynamics following the nutrient recovery, we first simplify the dynamics of the system. We find that at low methionine concentrations, $c_{\mathrm{met}} \approx 0$, the dynamics of the internal methionine concentration, Eq. (1) in the main text, can be approximated as

$$\left.\frac{\mathrm{d}c_{\mathrm{met}}}{\mathrm{d}t}\right|_{c_{\mathrm{met}}=0} = j_{\mathrm{in}} + h\phi_{\mathrm{MetA}}. \tag{S25}$$

Equation (S25) means that in the absence of methionine, there is no protein production, no drain to polyamine synthesis and, because there is no growth, there is no dilution. The dynamics is entirely determined by production $h\phi_{\mathrm{MetA}}$ and import $\tilde{j}_{\mathrm{in}}$. To get an estimate of the time-scale of this recovery, we calculate the rate change of methionine production

$$\left.\frac{\mathrm{d}\log c_{\mathrm{met}}}{\mathrm{d}t}\right|_{c_{\mathrm{met}}=0} = \frac{\tilde{j}_{\mathrm{in}} + h\phi_{\mathrm{MetA}}}{c_{\mathrm{met}}} \gg \frac{\tilde{j}_{\mathrm{in}} + h\phi_{\mathrm{MetA}}}{c_{\mathrm{met}}^{\mathrm{g}}} \approx 80\ \mathrm{h}^{-1}, \tag{S26}$$

where we used the upper bound that the methionine concentration $c_{\mathrm{met}}$ is lower than the growth steady state $c_{\mathrm{met}}^{\mathrm{g}} = 1.5 \cdot 10^{-4}$ M.

In comparison, the dynamics of the MetA abundance at $c_{\mathrm{met}} \approx 0$ is still mostly determined by degradation, as synthesis is small. We thus can estimate the rate change of the MetA abundance as

$$\left.\frac{\mathrm{d}\log\phi_{\mathrm{MetA}}}{\mathrm{d}t}\right|_{c_{\mathrm{met}}=0} = -\eta \approx -1.6\ \mathrm{h}^{-1}, \tag{S27}$$

When we compare Eq. (S26) with Eq. (S27), we see that methionine relaxes much faster than MetA. This common feature of cellular metabolism allows us to separate the dynamics of the system into a fast and slow phase, which we will use in the next section to calculate the unstable fixed-point.

#### 2. *Calculation of the unstable fixed-point*

The fact that the time scale of methionine recovery is faster than MetA means that, first, methionine concentration will increase, until consumption of methionine matches production. This is mathematically given by the nullcline $\mathrm{d}c_{\mathrm{met}}/\mathrm{d}t = 0$, Eq. (S8), at a constant $\phi_{\mathrm{MetA}}$ that equals the initial condition. Thereafter, both $c_{\mathrm{met}}$ and $\phi_{\mathrm{MetA}}$ will slowly increase or decrease together, until they finally converge to either the growth or dormancy steady state.

Whether they increase or decrease depends on whether the fast phase ends at a position above or below the unstable fixed-point on the methionine nullcline. A cell that started from the threshold $c_{\mathrm{met}} = 0$ & $\phi_{\mathrm{MetA}}^{\mathrm{th}}$ will land at the unstable fixed-point, $c_{\mathrm{met}} = c_{\mathrm{met}}^{\mathrm{unst}}$ & $\phi_{\mathrm{MetA}} = \phi_{\mathrm{MetA}}^{\mathrm{unst}}$. Because during this phase $\phi_{\mathrm{MetA}}$ is largely constant, we conclude that the threshold abundance of MetA, $\phi_{\mathrm{MetA}}^{\mathrm{th}}$, is roughly equal to the MetA abundance at the unstable fixed point,

$$\phi_{\mathrm{MetA}}^{\mathrm{th}} \approx \phi_{\mathrm{MetA}}^{\mathrm{unst}}. \tag{S28}$$

We can calculate the position of the unstable fixed point analogous to the above discussion of the two stable fixed points that led to the conditions of the three stability regimes, Eq. (S21-S23). In the regime around the unstable fixed point, drain has saturated, $u(K_{\mathrm{M}} \ll c_{\mathrm{met}}) = 1$. At the same time, $c_{\mathrm{met}}$ is still much smaller than at the growth fixed-point, $c_{\mathrm{met}}^{\mathrm{g}}$. This regime is given by $K_{\mathrm{M}} \ll c_{\mathrm{met}} \ll c_{\mathrm{met}}^{\mathrm{g}}$. In this case, the methionine nullcline is given by $\phi_{\mathrm{A}} \approx a_1 c_{\mathrm{met}} + a_2 - \tilde{j}_{\mathrm{in}}$ and the MetA abundance nullcline is given by $\phi_{\mathrm{MetA}} \approx \chi_{\mathrm{MetA}} \tilde{\eta}^{-1} c_{\mathrm{met}}$. Solving the MetA abundance nullcline for $c_{\mathrm{met}}$ and inserting it into the methionine nullcline gives us an estimate for the unstable fixed point,

$$\phi_{\mathrm{MetA}}^{\mathrm{unst}} = \frac{a_2 - \tilde{j}_{\mathrm{in}}}{1 - a_1 \tilde{\eta} \chi_{\mathrm{MetA}}^{-1}}, \tag{S29}$$

or written in the original parameters

$$\phi_{\mathrm{MetA}}^{\mathrm{unst}} = \frac{1}{h} \frac{k_{\mathrm{d}} - j_{\mathrm{in}}}{1 - h c_{\mathrm{met}}^{\mathrm{pro}} \eta \chi_{\mathrm{MetA}}^{-1}}. \tag{S30}$$

Equation (S30) then defines the growth threshold, because of Eq. (S28).

#### *C. Calculation of lag time*

In this section, we aim to estimate the lag times of the population. Lag times are caused by the recovery of a small, growing subpopulation. For this reason, in this section, we first study the heterogeneity of gene expression that leads to the subpopulations, second, what determines the size of the subpopulations and third, how the size of the subpopulations affects lag time.

##### *1. Heterogeneity in MetA abundance*

We investigate how the existence of a threshold can lead to a separation of the population into two growth and dormancy states. Generally, it is known that gene expression is a highly heterogeneous process, due to the stochastic synthesis and degradation of mRNAs and proteins [10,11]. Phenomenologically, the abundance of proteins such as MetA is well described by a gamma distribution [10,11] ,

$$p(\phi_{\mathrm{MetA}}) = \frac{{\phi_{\mathrm{MetA}}}^{a-1}\exp(-\phi_{\mathrm{MetA}}b^{-1})}{\Gamma(a)b^{a}}, \tag{S31}$$

where $a$ is the number of mRNA synthesized per cell cycle and $b$ the number of proteins synthesized per mRNA. Due to the nature of the gamma distribution, $a$ also equals the inverse of the noise, $\sqrt{a}^{-1} = \sigma/\xi$ and b equals the Fano factor, $b = \sigma^2/\xi$, where $\sigma$ is the standard deviation and $\xi = ab$ is the average. In Table S2, we estimate the average MetA abundance, $\phi_{\mathrm{A}}^{*} = 6.13 \cdot 10^{-4} = \xi$. In order to estimate the noise, we take the median noise, $\sqrt{a}^{-1}{}_{\mathrm{median}} = \sigma/\xi_{\ \mathrm{median}}$, of proteins with an average copy number of more than 10 [11] and find $\sqrt{a}^{-1}{}_{\mathrm{median}} = 0.4$. We use this value to plot the gamma distribution of MetA in the main text. However, we want to note that neither of these values appear in the theoretical predictions of the lag time, calculated further down in this Supplementary Information in Eq. (S40).

#### 2. *Calculation of lag times*

During starvation, the abundance of MetA decays exponentially due to thermal inactivation [8],

$$\phi_{\mathrm{MetA}}(t) = \phi_{\mathrm{MetA}}(0)\exp(-\eta t), \tag{S32}$$

and the MetA abundance in many cells will falls below the threshold $\phi_{\mathrm{MetA}}^{\mathrm{th}}$ required for growth, calculated in Eq. (S30). By saying that all cells above the threshold will recover and cells below the threshold will not recover, we implicitly neglect gene expression noise during recovery.

We can calculate the number of recovering cells by integrating the probability distribution, Eq. (S31), obtained after a starvation time $\tau$, from the growth threshold $\phi_{\mathrm{MetA}}^{\mathrm{th}}$ to infinity,

$$\Pi\left(\phi_{\mathrm{MetA}}^{\mathrm{th}}, \tau\right) = \int_{\phi_{\mathrm{MetA}}^{\mathrm{th}}}^{\infty} p\left(\phi_{\mathrm{MetA}}(\tau)\right) \mathrm{d}\phi_{\mathrm{MetA}}. \tag{S33}$$

We can estimate the resulting lag time by assuming that recovery of individual cells is fast. Then, lag time will be entirely determined by a growing population $\Pi$ (growing immediately at growth rate $\mu$) and a dormant population $1-\Pi$ (that will never grow). These dynamics of the two subpopulations and the resulting lag time is sketched in the main text in Figure 5.

The resulting biomass growth of the population, is given by

$$M(t) = M(0)\left(\left(1 - \Pi\left(\phi_{\mathrm{MetA}}^{\mathrm{th}}, \tau\right)\right) + \Pi\left(\phi_{\mathrm{MetA}}^{\mathrm{th}}, \tau\right)\exp(\mu t)\right). \tag{S34}$$

Note that in this description we are focusing on describing growth, and we are neglecting the slow decay of optical density observed during starvation and lag, see Fig. 1 of the main text.

In order to calculate the lag time, we study the asymptotic growth, when the growing subpopulation has out-grown the non-growing population. In this regime, $\Pi\left(\phi_{\mathrm{MetA}}^{\mathrm{th}}, \tau\right)\exp(\mu t) \gg 1$, and the biomass growth reaches

$$M(t) = M(0)\Pi\left(\phi_{\mathrm{MetA}}^{\mathrm{th}}, \tau\right)\exp(\mu t) = M(0)\exp\left(\mu\left(t - T_{\mathrm{lag}}\right)\right) \quad \text{(S35)}$$

with the lag time

$$T_{\mathrm{lag}} = \frac{1}{\mu}\ln\left(\frac{1}{\Pi\left(\phi_{\mathrm{MetA}}^{\mathrm{th}}, \tau\right)}\right). \quad \text{(S36)}$$

#### 3. *Scaling of lag time with physiological parameters*

To understand the impact of experimental perturbations, such as these performed in the main text, we are looking for analytical scaling laws between the measurable output lag time and biological parameters such as MetA pre-expression, starvation time and degradation rate. For this, we note that long lag times are caused by very small growing subpopulations. The growing subpopulation $\Pi\left(\phi_{\mathrm{MetA}}^{\mathrm{th}}, \tau\right)$ becomes small when the threshold $\phi_{\mathrm{MetA}}^{\mathrm{th}}$ is much larger than the mean of the distribution $p\left(\phi_{\mathrm{MetA}}(\tau)\right)$. Thus, the growing subpopulation is represented by the tail of the gamma distribution, which is exponential with MetA abundance $\phi_{\mathrm{MetA}}(t)$ that decreases during starvation, and is given by

$$p(\phi_{\mathrm{MetA}}) \propto \exp(-\phi_{\mathrm{MetA}}(t)b^{-1}). \quad \text{(S37)}$$

We can then obtain the growing subpopulation after a certain starvation time $\tau$ using Eq. (S33),

$$\Pi\left(\phi_{\mathrm{MetA}}^{\mathrm{th}}, \tau\right) \propto \exp\left(-\phi_{\mathrm{MetA}}^{\mathrm{th}} b^{-1}\exp(\eta\tau)\right). \quad \text{(S38)}$$

The growing subpopulation thus scales with a

(1) double-exponential form with starvation time $\tau$,
(2) double-exponential form with degradation rate $\eta$,
(3) exponentially with the growth threshold $\phi_{\mathrm{MetA}}^{\mathrm{th}}$,
(4) exponentially with the inverse of the mean pre-expression $\xi = ab$.

By plugging in the growing subpopulation, Eq. (S38), our derivation of lag time Eq. (S36), we find that lag time scales proportional to,

$$T_{\text{lag}} \propto \text{const} + \phi_{\text{MetA}}^{\text{th}} b^{-1} \exp(\eta\tau). \tag{S39}$$

In summary, lag time scales approximately

(1) exponentially with starvation time $\tau$,
(2) exponentially with degradation rate $\eta$,
(3) linearly with the growth threshold $\phi_{\text{MetA}}^{\text{th}}$,
(4) linearly with the inverse of the mean pre-expression $\xi = ab$.

#### *D. Estimates of physiological parameters*

Using published physiological characterization of *E. coli,* we can estimate the parameters used in the theory. The results are summarized in Table S2.

##### *1. Growth physiology*

At 45 degrees, *E. coli* in minimal medium supplemented with glycerol has a growth rate of about $\mu^{\mathrm{g}} = 0.45\ \mathrm{h}^{-1}$, see main text. The typical cell volume in minimal medium is about $V = 1.3\ \mathrm{\mu m}^3 = 1.3 \cdot 10^{-15}\ \mathrm{l}$, with a protein mass of about $M = 165\ \mathrm{pg}$ [6]. The exact numbers of volume and mass do not matter for the bistability conditions, but serve the purpose to estimate parameters such as the MetA copy number per cell.

##### *2. Methionine and related compounds*

The internal concentration of free methionine (molar mass $m_{\mathrm{met}} = 149\ \mathrm{g\ mol}^{-1}$) in cells growing in minimal medium is about $c_{\mathrm{met}}^{\mathrm{g}} = 150\ \mathrm{\mu M}$ at 37ºC [12]. Using that the internal methionine concentration is not much different at 45ºC, we estimate the proportionality constant between methionine concentration and growth rate, $k_{\mathrm{p}} = \mu^{\mathrm{g}} {c_{\mathrm{met}}^{\mathrm{g}}}^{-1} = 3.0 \cdot 10^3\ \mathrm{M\ h}^{-1}$.

The spermidine concentration, the major polyamine that drains methionine, is $c_{\mathrm{spe}} = 1.4\ \mathrm{mM}$ at 37ºC [6]. The synthesis of each spermidine requires one S-adenosyl-methionine, which is directly draining the methionine pool. The resulting byproduct, S-methyl-thioadenosine cannot be recycled back to methionine without passage through MetA.

While the spermidine pool is 10-fold larger than the free methionine pool, we note that the majority of the methionine is bound inside proteins, where they make up about 3.23% of the protein mass, see Table S2 and S3. This corresponds to a methionine concentration of about $c_{\mathrm{met}}^{\mathrm{pro}} = 3.23\%\ M m_{\mathrm{met}}^{-1} V^{-1} = 27.5\ \mathrm{mM}$.

Thus, we conclude that during growth, about $27.5\ \mathrm{mM}/1.4\ \mathrm{mM} = 20$ fold more methionine goes into protein synthesis than into spermidine synthesis. In terms of systems parameters, this ratio can be written as

$$\frac{k_\mathrm{d}}{\mu^\mathrm{g} c_\mathrm{met}^\mathrm{pro}} = \frac{1}{20}, \tag{S40}$$

when we assume that $u(c_\mathrm{met})$ is saturated at $u(c_\mathrm{met}) = 1$ at full growth.

##### 3. *Enzymatic parameters of MetA*

From the measured copy number of MetA, $N_\mathrm{MetA} = 1711$ (Li et al., 2014), we can estimate the abundance $\phi_\mathrm{MetA} = N_\mathrm{MetA} m_\mathrm{MetA} M^{-1}\, N_\mathrm{A}^{-1} = 6.13 \cdot 10^{-4}$, if we use the Avogadro number, $N_\mathrm{A} = 6.022 \cdot 10^{23}\mathrm{mol}^{-1}$, the typical cell mass $M = 165$ pg and the molar mass of MetA, $m_\mathrm{MetA} = 3.56 \cdot 10^4\ \mathrm{g\ mol}^{-1}$. From the MetA abundance, we estimate the methionine production rate by MetA, $h$, as $h = (c_\mathrm{met} +\ c_\mathrm{met}^\mathrm{pro})\mu^\mathrm{g} \phi_\mathrm{MetA}^{\mathrm{g}\ -1} = 20.3\ \mathrm{Mh}^{-1}$.

The degradation rate of MetA is about $\eta(37°\mathrm{C}) = 1.1\ \mathrm{h}^{-1}$ at 37°C and increases to $\eta(45°\mathrm{C}) = 1.6\ \mathrm{h}^{-1}$ at 45°C [8]. In order to maintain a steady state MetA abundance of $\phi_\mathrm{MetA}^\mathrm{g} = 2.6 \cdot 10^{-5}$ at growth rate $\mu^\mathrm{g} = 0.45\ \mathrm{h}^{-1}$, the synthesis fraction of MetA, defined as the fraction of the protein mass synthesis that is directed to MetA, has to be $\chi_\mathrm{MetA} = \phi_\mathrm{MetA}\, (\eta + \mu^\mathrm{g})\, \mu^{\mathrm{g}^{-1}} = 2.79 \cdot 10^{-3}$, where we used Eq. (S11).

##### 4. *Compound parameters*

The above parameters together determine the essential parameters of the system, $a_1$, $a_2$, $a_3$ and $\tilde{\eta}$ that are summarized in Table S2.

### II. SUPPLEMENTARY FIGURES AND TABLES

#### A. Supplementary Figures

```
AFABRUM           MPIKIPDTLPAFETLVHEGVRVMTETAAIRQDIRPLQIGLLNLMPNKIKTEIQMARLVGA  60
PPOLYMYXA         MPIKIPDTLPAKEVLEGENIFVMDESLAYHQDIRPLRIAILNLMPTKETTETQLLRLVGN  60
KAERGOGENES       MPIRVQDELPAVNFLRDENVFVMTTSRATTQEIRPLKVLILNLMPKKIETENQFLRLLSN  60
ECOLI             MPIRVPDELPAVNFLREENVFVMTTSRASGQEIRPLKVLILNLMPKKIETENQFLRLLSN  60
STYPHIMURIUM      MPIRVLDELPAVNFLREENVFVMTTSRASGQEIRPLKVLILNLMPKKIETENQFLRLLSN  60
                  ***:: * *** : *  *.: **  : *  *:****:: :*****.*  ** *: **:.

                                                    Q96K
AFABRUM           SPLQVELSLIRIGGHRAKNTPEEHLLSFYQTWEEVRHRKFDGFIITGAPIELLDYEDVTY  120
PPOLYMYXA         TPLQVDVTLVHMKSHVSKNTSQEYLNMFYKTFDEIKNSRFDGMVITGAPVEQLEFEDVNY  120
KAERGOGENES       SPLQVDIQLLRIDARESRNTPTEHLNNFYCNFEDIREQNFDGLIVTGAPLGLVEFNDVAY  120
ECOLI             SPLQVDIQLLRIDSRESRNTPAEHLNNFYCNFEDIQDQNFDGLIVTGAPLGLVEFNDVAY  120
STYPHIMURIUM      SPLQVDIQLLRIDARESRNTPAEHLNNFYCNFDDICDQNFDGLIVTGAPLGLVEFNDVAY  120
                  :****:: *::: .: ::**  *:*  ** .:::: . .***::****:  ::::** *

                     I124L
AFABRUM           WNEMQQIFEWTQTNVHSTLNVCWGAMAAIYHFHGVPKYELKEKAFGVYRHRNLSPSSIYL  180
PPOLYMYXA         WEEIRQIFEWTKTNVTSTMHICWASQAGLYHHFDVPKYGLDTKCFGVFPHTVIKPKVKLL  180
KAERGOGENES       WPQIKQVLEWAKDHVTSTLFVCWAVQAALNILYGIPKQTRTEKISGVYEHHILHPHALLT  180
ECOLI             WPQIKQVLEWSKDHVTSTLFVCWAVQAALNILYGIPKQTRTEKLSGVYEHHILHPHALLT  180
STYPHIMURIUM      WPQIRQVLEWAKDHVTSTLFVCWAVQAALNILYGIPKQTRTDKLSGVYEHHILHPHALLT  180
                  * :::*::**:: :* **: :**.  *.:   ..:**     *  **: *  : *

                                                                 I229T/Y
AFABRUM           NGFSDDFQVPVSRWTEVRRADIEKHPELEILMESDEMGVCLAHEKAGNRLYMFNHVEYDS  240
PPOLYMYXA         RGFDELFYAPHSRHTEVRREDIDHIAELEVLSESEEAGVYLVATLDGKQIFVTGHSEYDP  240
KAERGOGENES       RGFDDTFLAPHSRYADFPAGLIRDYTDLDILAETEDGDAYLFASKDKRIAFVTGHPEYDP  240
ECOLI             RGFDDSFLAPHSRYADFPAALIRDYTDLEILAETEEGDAYLFASKDKRIAFVTGHPEYDA  240
STYPHIMURIUM      RGFDDSFLAPHSRYADFPAALIRDYTDLEILAETEEGDAYLFASKDKRIAFVTGHPEYDA  240
                  .**.: * .* ** ::.    * .  :*::* *::: .. *       .  :: .* ***

                       F247Y                 N267D
AFABRUM           TSLADEYFRDVNSGVPIKLPHDYFPHNDPELAPLNRWRSHAHLFFGNWINE-IYQTTPYD  299
PPOLYMYXA         LSLKWEYDRDVAKGLDIDVPKNYFPNDDPERIPPSIWRAHANLLFSNWLNYYVYQETPYD  300
KAERGOGENES       HTLASEYFRDLEAGLAPQLPDNYFPNNDPNNKPRATWRSHGNLLFANWLNYYVYQITPYD  300
ECOLI             QTLAQEFFRDVEAGLDPDVPYNYFPHNDPQNTPRASWRSHGNLLFTNWLNYYVYQITPYD  300
STYPHIMURIUM      HTLAGEYFRDVEAGLNPEVPYNYFPKNDPQNIPRATWRSHGNLLFTNWLNYYVYQITPYD  300
                   :*  *: **:  *:  .:* :***::**:  *   **:*.:*:* **:*  :** ****

AFABRUM           PQAIGKLAA      308
PPOLYMYXA         IGPQI----      305
KAERGOGENES       LRHMNPTLD      309
ECOLI             LRHMNPTLD      309
STYPHIMURIUM      LRHMNPTLD      309
```

**Figure S1. Clustal Omega (1.2.4)[13] multiple sequence alignment (MSA) of MetA protein sequences of mesophilic bacteria with known thermolabile MetA[2,14–17].** MSA of *Agrobacterum fabrum* C58, *Paenibacillus polymyxa* SC2, *Klebsiella aerogenes* ATCC 13048,

*Escherichia coli* K12 MG1655 and *Salmonella Typhimurium* LT2 ATCC 700720. Highlighted residues indicate mutations known to increase thermal tolerance, identified by Mordukhova et al: Q96K, I124L, 1229T, I229Y, F247Y and N267D [8,18]. Consensus symbols ‘*’, ‘:’ and ‘.’ indicate complete conservation, conservation of groups of strongly similar physicochemical properties, and conservation of groups of weakly similar physicochemical properties respectively. MetA of *P. polymyxa* and *A. fabrum* have significantly diverged from *E. coli* and include several mutations identified by Mordukhova to stabilize MetA, including Q96K, F247Y and N267D. Despite these mutations, MetA of *P. polymyxa* and *A. fabrum* remains thermolabile [15,17].

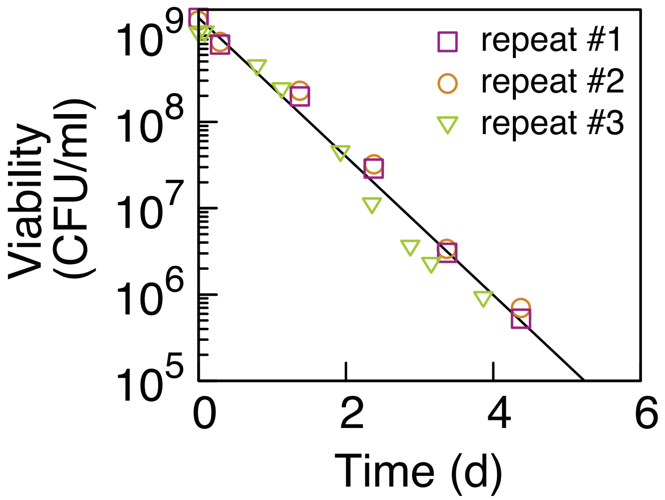

**Figure S2. Survival kinetics of *E. coli* K-12 during starvation at 45ºC.** Viability of *E. coli*, measured in colony forming units on LB agar plates incubated at 37ºC over the course of 5 days. Viability decreases exponentially at a rate of 0.081/h. After 5 h of starvation, viability has decreased to about $\exp(5\ \mathrm{h} \cdot 0.081\ h^{-1}) = 67\%$ . Death rate at 45ºC is about 5-fold faster than at 37ºC: 0.081/h compared to 0.018/h [19].

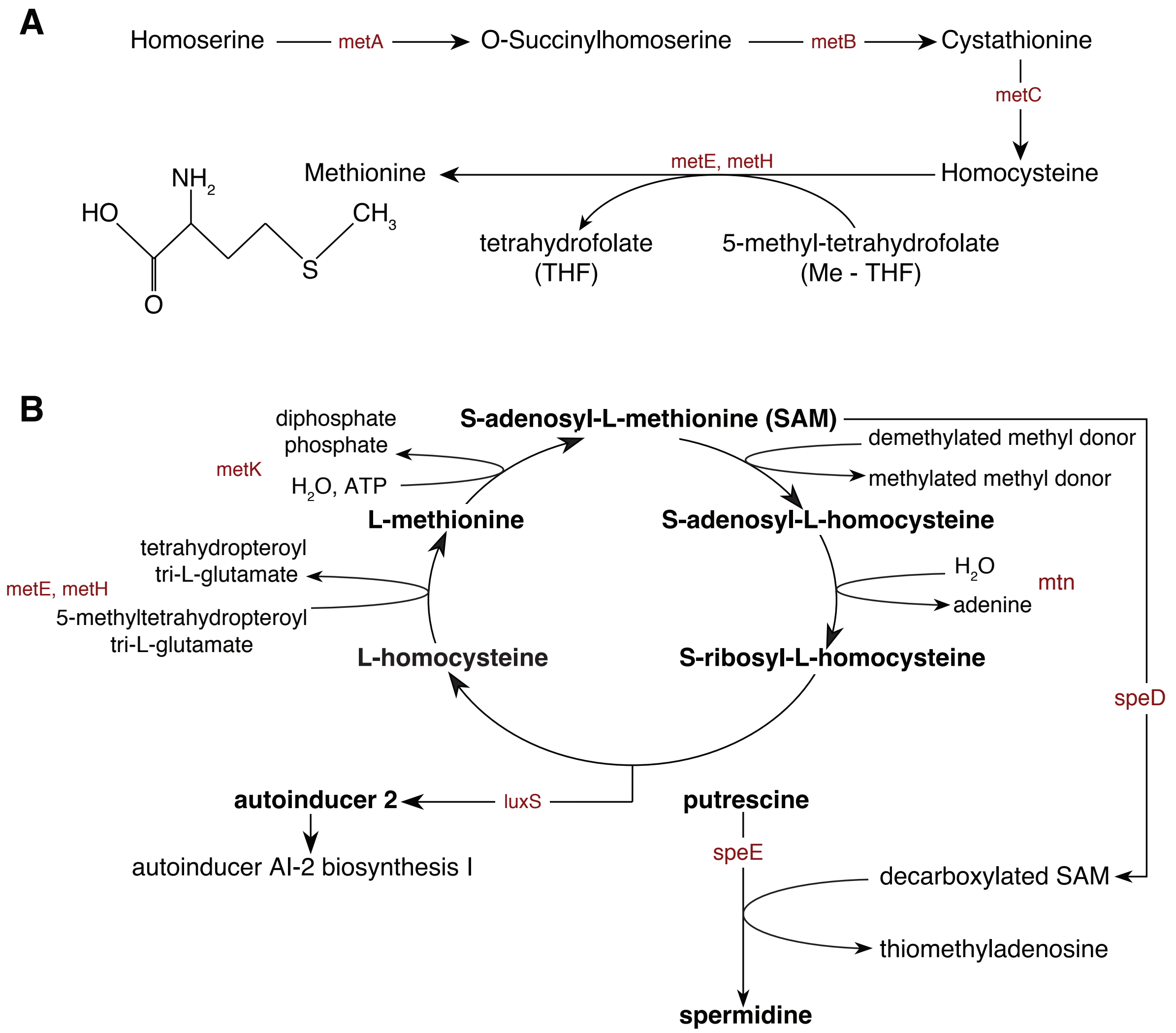

**Figure S3. L-methionine biosynthetic pathway and SAM cycle in E. coli K-12. (A)** Methionine biosynthesis pathway. The genes encoding the enzymes that catalyze each step are indicated in red: *metA* codes for homoserine O-succinyltransferase, *metB* for cystathionine $\gamma$-synthase, *metC* for cystathionine $\beta$-lyase, *metH* or *metE* for methionine synthase. The chemical structure of L-methionine is also reported. It contains an amino group (in the $NH_3^+$ form under biological conditions), an $\alpha$-carboxylic acid group (in the -COO form under biological conditions) and an S methyl thioether side chain. **(B)** S-adenosyl-L-methionine cycle I. Methionine conversion into SAM, the major methyl donor in the cell. As in panel A, the genes encoding the enzymes that catalyze each step are indicated in red. Methionine is activated by condensation with ATP to form

S-adenosyl-L-methionine (AdoMet or SAM). The synthesis of AdoMet is catalyzed by methionine adenosyltransferase, the *metK* gene product. Then, SAM donates its methyl group and it is converted to S-adenosyl-L-homocysteine (SAH). SAH is first hydrolyzed to S-ribosyl-L-homocysteine by 5'-methylthioadenosine, followed by conversion to L-homocysteine by S-ribosylhomocysteine lyase. The cycle continues with the methylation of L-homocysteine to L-methionine using a methyl group from a methylated folate. Finally, the cycle is completed with the regeneration of SAM by methionine adenosyltransferase. Within the cycle, a fraction of methionine that is not recycled, because SAM is used to produce polyamines, such as spermidine. Polyamines are polycations involved in many biological processes including binding to nucleic acids, stabilizing membranes and stimulation of enzymes essentials for growth. They are also often used in responses to stress such as heat stress [5]. In this case, the enzyme SAM decarboxylase (from gene *speD*) converts SAM into decarboxylated SAM, which acts as a cofactor for spermidine biosynthesis. Then, spermidine synthase (from gene *speE*) converts putrescine to spermidine in the presence of decarboxylated SAM. The autoinducer AI-2 is an *E. coli* signaling molecule and could be used for cell-cell communication (e.g. quorum sensing), allowing bacterial populations to coordinate gene expression as a function of cell density. Its biosynthesis is catalyzed by the enzyme S-ribosylhomocysteine lyase, from the gene *luxS*, which uses cleavage of S-ribosylhomocysteine. The image is based on information acquired from [20,21].

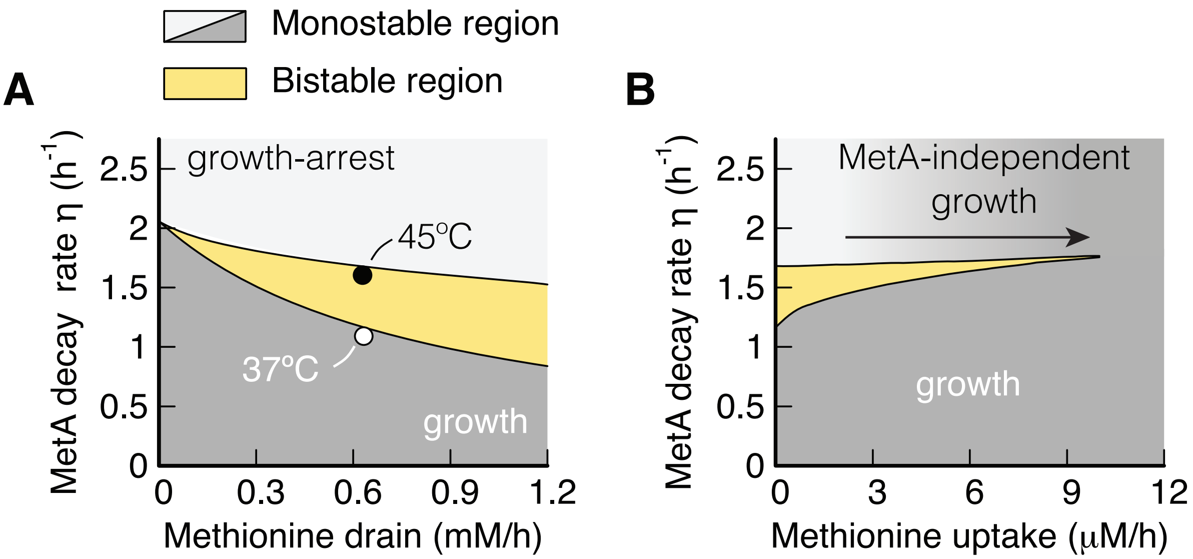

**Figure S4. Stability analysis as function of MetA decay rate, methionine drain and methionine uptake. (A)** Stability diagram showing monostable regimes, growing (dark grey region) non-growing (light grey region), and the bistable regime (yellow region) as function of the parameters MetA degradation rate and methionine drain, without methionine influx. Black & white symbols: estimates of values at 37 °C and 45 °C, see Table S2. Methionine drain is essential for the observed bistability, as it proves the mathematically required non-linearity [1].**(B)** Stability diagram as function of the MetA degradation rate and uptake rate of external methionine, for fixed methionine drain $j_d = 0.6\ \mathrm{mM\ h^{-1}}$. The shaded region (light grey to dark grey) indicates that bacteria are increasingly able to grow on imported methionine, i.e. independent of MetA.

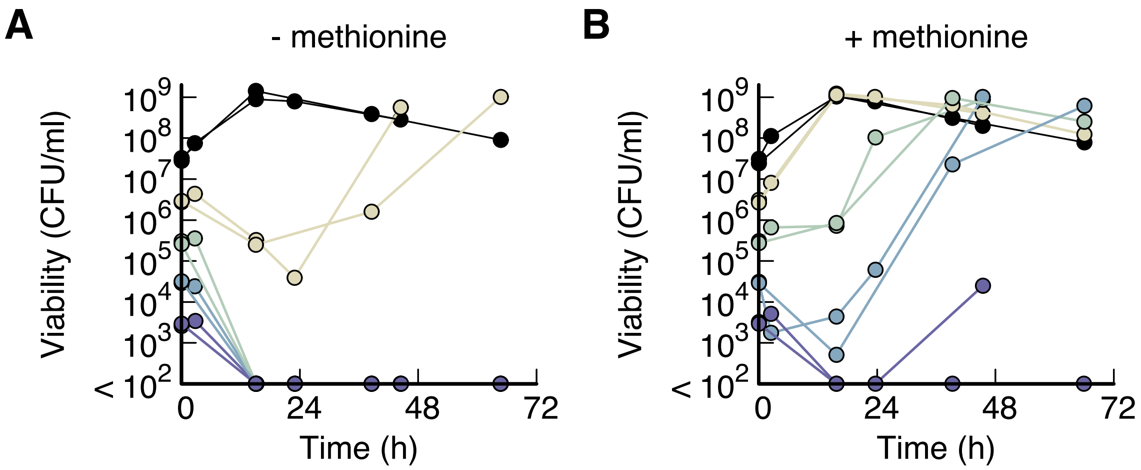

**Figure S5. Regrowth of *E. coli* following dilutions to various initial population densities, with and without methionine. (A)** Growth of *E. coli* diluted during exponential growth in minimal medium with glycerol (20 mM) at 44ºC, to varying cell densities. Only cultures with initial cell densities exceeding $10^6$ CFU/mL exhibited regrowth within 72 hours (yellow and black curves). **(B)** Regrowth of *E. coli* under the same conditions as in (A), but with the addition of methionine (0.067 mM). All cultures with initial viabilities larger than $10^4$ CFU/mL regrew within 72 hours.

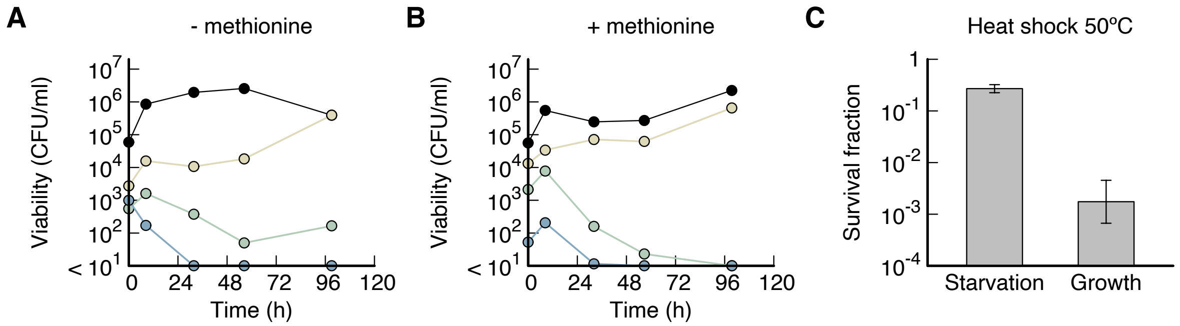

**Figure S6. Yeast. (A-B)** Growth of *S. cerevisiae* diluted during exponential growth in NBM with 0.5% glucose to different cell densities, (A) with and (B) without the addition of methionine (0.067mM). Only $10^1$ (black) and $10^2$-fold (yellow) dilutions exhibited regrowth within 96 hours, regardless of the presence of methionine. **(C)** Killing of *S. cerevisiae* by heat shock (10 minutes at 50ºC), both during growth and following 2 h starvation at 39ºC. Growth arrest due to carbon starvation aided the survival of *S. cerevisiae* during heat shocks.

#### *B. Supplementary Tables*

##### *1. Table S1.*

***Growth rates with and without methionine at different temperatures.*** *Each value is the average of three independent repeats with a typical standard deviation of 0.01.*

| Temperature (ºC) | Growth rate w/o met. (1/h) | Growth rate w/ met. (1/h) |
|---|---|---|
| 25 | 0.30 | 0.33 |
| 27 | 0.38 | 0.42 |
| 29 | 0.48 | 0.53 |
| 31 | 0.54 | 0.60 |
| 33 | 0.60 | 0.69 |
| 35 | 0.65 | 0.78 |
| 37 | 0.70 | 0.86 |
| 39 | 0.75 | 0.95 |
| 40 | 0.78 | 0.97 |
| 41 | 0.75 | 0.96 |
| 42 | 0.71 | 0.95 |
| 43 | 0.62 | 0.81 |
| 44 | 0.54 | 0.72 |
| 45 | 0.45 | 0.62 |
| 46 | 0 | 0.49 |
| 47 | 0 | 0.14 |
| 48 | 0 | 0 |

#### 2. *Table S2.*

***Summary of system parameters.*** *Top: Physiological parameters from this work and the literature. If measurements at 45°C were not available, we used values obtained at 37°C. Note that the theoretical predictions tested in Fig. 6 are independent of the physiological parameters. The parameters presented here serve solely an illustrative purpose by estimating sizes of the described effects. Middle: Derived estimates for parameters used to illustrate Figure 3 in the main text. Bottom: Rescaled parameters used in the theory.*

##### Physiological parameters

| **Name** | **Symbol** | **Value** | **Unit** | **Reference** | **Comment** |
|---|---|---|---|---|---|
| Growth rate | $\mu^{\mathrm{g}}$ | 0.45 | 1/h | This work. | *E. coli* K-12, glycerol minimal medium, 45°C |
| MetA copy number | $N_{\mathrm{MetA}}$ | 1711 | - | Ref. [22] | *E. coli* K-12 at 37 °C, glucose minimal medium |
| Avogadro constant | $N_{\mathrm{A}}$ | $6.02 \cdot 10^{23}$ | 1/mol | - | - |
| Typical internal methionine concentration | $c_{\mathrm{met}}^{\mathrm{g}}$ | $1.5 \cdot 10^{-4}$ | M | Ref. [12] | *E. coli* B/r at 37 °C, in glucose minimal medium |
| Protein dry mass of *E. coli* | $M$ | $165 \cdot 10^{-15}$ | g | Ref. [6] | *E. coli* B/r at 37 °C, in glucose minimal medium |
| Volume of *E. coli* | $V$ | $1.30 \cdot 10^{-15}$ | l | Ref. [6] | *E. coli* B/r at 37 °C, in glucose minimal medium |
| MetA decay rate | $\eta$ | 37°C: 1.1<br>45°C: 1.6 | 1/h | Ref. [8] | *E. coli* JW3973, in glucose minimal medium |
| MetA molar mass | $m_{\mathrm{MetA}}$ | $3.56 \cdot 10^{4}$ | g/mol | | |
| Spermidine concentration | $n_{\mathrm{spe}}$ | $1.10 \cdot 10^{6}$ | 1/cell | Ref. [6] | *E. coli* B/r at 37 °C, in glucose minimal medium |
| Methionine affinity constant | $K_{\mathrm{M}}$ | 10 | µM | Estimated in this work. | - |

##### Model parameters

| **Name** | **Symbol** | **Value** | **Unit** | **Calculation** | **Comment** |
|---|---|---|---|---|---|
| Rate constant of methionine dependent growth | $k_{\mathrm{p}}$ | $3.0 \cdot 10^{3}$ | 1/(M h) | $k_p = \mu^{\mathrm{g}} {c_{\mathrm{met}}^{\mathrm{g}}}^{-1}$ | Converts internal methionine concentration into growth. |
| Methionine concentration<br>(in proteins) | $c_{\mathrm{met}}^{\mathrm{pro}}$ | $2.75 \cdot 10^{-2}$ | M | $c_{\mathrm{met}}^{\mathrm{pro}} = 3.23\% \frac{M}{V} / 149 \frac{\mathrm{g}}{\mathrm{mol}}$ | 3.23% is the fraction of the protein mass that is methionine and 149 g/mol is the molar mass of methionine. Both parameters are from Table S3. |
| Spermidine concentration | $c_{\mathrm{spe}}$ | $1.41 \cdot 10^{-3}$ | M | $c_{\mathrm{spe}} = n_{\mathrm{spe}} N_{\mathrm{A}} / V$ | Per spermidine one methionine is used. |

| | | | | | |
|---|---|---|---|---|---|
| Active MetA mass fraction | $\phi^{\mathrm{g}}_{\mathrm{MetA}}$ | $6.13 \cdot 10^{-4}$ | - | $\phi_{\mathrm{MetA}} = \dfrac{N_{\mathrm{MetA}} m_{\mathrm{MetA}}}{M\, N_{\mathrm{A}}}$ | Does not include misfolded or degraded MetA. |
| MetA synthesis fraction | $\chi_{\mathrm{MetA}}$ | $2.79 \cdot 10^{-3}$ | | $\chi_{\mathrm{MetA}} = \phi^{\mathrm{g}}_{\mathrm{MetA}} \dfrac{(\eta + \mu^{\mathrm{g}})}{\mu^{\mathrm{g}}}$ | Estimated for 45ºC. |
| Methionine production rate by MetA | $h$ | 20.3 | M/h | $h = \dfrac{\left(c^{\mathrm{pro}}_{\mathrm{met}} + c^{\mathrm{g}}_{\mathrm{met}}\right) \mu^{\mathrm{g}}}{\phi^{*}_{\mathrm{A}}}$ | - |
| Strength of the methionine drain | $k_{\mathrm{d}}$ | $6.3 \cdot 10^{-4}$ | M/h | $k_{\mathrm{d}} = c_{\mathrm{spe}} \mu^{\mathrm{g}}$ | - |

#### Compound parameters

| **Name** | **Sym bol** | **Value** | **Unit** | **Calculation** | **Comment** |
|---|---|---|---|---|---|
| Rescaled protein synthesis | $a_1$ | 4.06 | 1/M | $k_{\mathrm{p}} c^{\mathrm{pro}}_{\mathrm{met}} h^{-1}$ | - |
| Rescaled methionine drain | $a_2$ | $3.1 \cdot 10^{-5}$ | - | $k_{\mathrm{d}} h^{-1}$ | - |
| Rescaled dilution term | $a_3$ | 148 | $1/\mathrm{M}^2$ | $k_{\mathrm{p}} h^{-1}$ | - |
| Rescaled decay rate | $\tilde{\eta}$ | 37°C: $0.37 \cdot 10^{-3}$<br>45°C: $0.53 \cdot 10^{-3}$ | M | $\eta k_p^{-1}$ | Temperature dependent. |

#### 3. *Table S3.*

*Amino acid frequency, and relative abundance of amino acids, in percent of total protein mass, estimated based on the codon frequency of E. coli from: http://www.kazusa.or.jp/codon/*

| **Amino acid** | **Frequency** | **Molar-mass (g/mol)** | **Relative abundance** |
|---|---|---|---|
| Glycine | 7.34% | 75.07 | 4.28% |
| Glutamate | 5.75% | 147.13 | 6.58% |
| Aspartate | 5.14% | 133.1 | 5.32% |
| Valine | 7.09% | 117.15 | 6.45% |
| Alanine | 9.51% | 89.09 | 6.58% |
| Arginine | 5.54% | 174.2 | 7.49% |
| Lysine | 4.40% | 146.19 | 4.99% |
| Asparagine | 3.94% | 132.12 | 4.04% |
| Methionine | 2.79% | 149.21 | 3.23% |
| Isoleucine | 5.99% | 131.18 | 6.10% |
| Threonine | 5.39% | 119.12 | 4.98% |
| Tryptophan | 1.54% | 204.23 | 2.44% |
| Cysteine | 1.17% | 121.15 | 1.10% |
| Tyrosine | 2.87% | 181.19 | 4.04% |
| Phenylalanine | 3.91% | 165.19 | 5.02% |
| Serine | 5.81% | 105.09 | 4.74% |
| Glutamine | 4.44% | 146.15 | 5.04% |
| Histidine | 2.27% | 155.16 | 2.73% |
| Leucine | 10.67% | 131.18 | 10.87% |
| Proline | 4.45% | 115.13 | 3.98% |
